# Generation of a transgenic cephalopod

**DOI:** 10.64898/2026.09.23.753964

**Authors:** Tessa G. Montague, Connor J. Gibbons, G. Thomas Barlow, Elizaveta V. Bashkirova, Yilan Xu, Eva C. Castagna, Thomas J. Mungioli, Kelly A. Patrick, Adriana Nemes, Aalok Varma, Loren L. Looger, Caroline B. Albertin, Stavros Lomvardas, Richard Axel

## Abstract

Coleoid cephalopods (cuttlefish, octopus, and squid) are marine mollusks with elaborate nervous systems that support a diverse repertoire of complex behaviors. These include the neural control of the color, pattern, and texture of the skin, facilitating both adaptive camouflage and innate patterning that may reflect internal state. The development of transgenic cephalopods expressing fluorescent proteins, optogenetic actuators, and reporters of neural activity would contribute a new and important technology to cephalopod biology. The generation of transgenic cephalopods, however, has remained a major challenge. Here, we report the development of stable transgenic dwarf cuttlefish (*Ascarosepion bandense*) expressing ubiquitous nuclear-localized mScarlet, a red fluorescent protein. We evaluated multiple strategies for transgenesis, and established cuttlefish lines using both CRISPR and the transposons Sleeping Beauty and Minos. The stable expression of transgenes enabled live imaging of cell dynamics during embryonic development. The Minos transposon emerged as the most efficient transgenesis strategy and is adaptable to promoters and transgenes of choice. These strategies now enable the generation of diverse genetic tools for mechanistic studies of cephalopod biology.

## Introduction

Coleoid cephalopods (cuttlefish, octopus, and squid) shared a common ancestor with vertebrates ∼600 million years ago^1^ (Figure 1A) yet independently evolved large brains and a sophisticated behavioral repertoire that includes episodic-like memory^2^, problem-solving^3^, adaptive camouflage^4^ (Figure 1B), and complex social behaviors^5^ (Figure 1C). During camouflage, cephalopods dynamically alter the color, pattern, and texture of their skin to create an approximation of their surroundings^4^. The cephalopod nervous system transforms the visual world into adaptive skin patterns that are generated using chromatophores, neuromuscular pigment sacs that expand and contract under the control of motor neurons projecting from the brain, permitting color change in milliseconds^6^ (Figures 1D,E). In addition to camouflage, cephalopods create neurally-controlled skin displays during social encounters, exhibiting sequences of stereotyped skin patterns that may communicate internal states^5^ (Figure 1C). Thus, the skin of cephalopods serves as a dynamic external display of internal neural activity.

**Figure 1.**
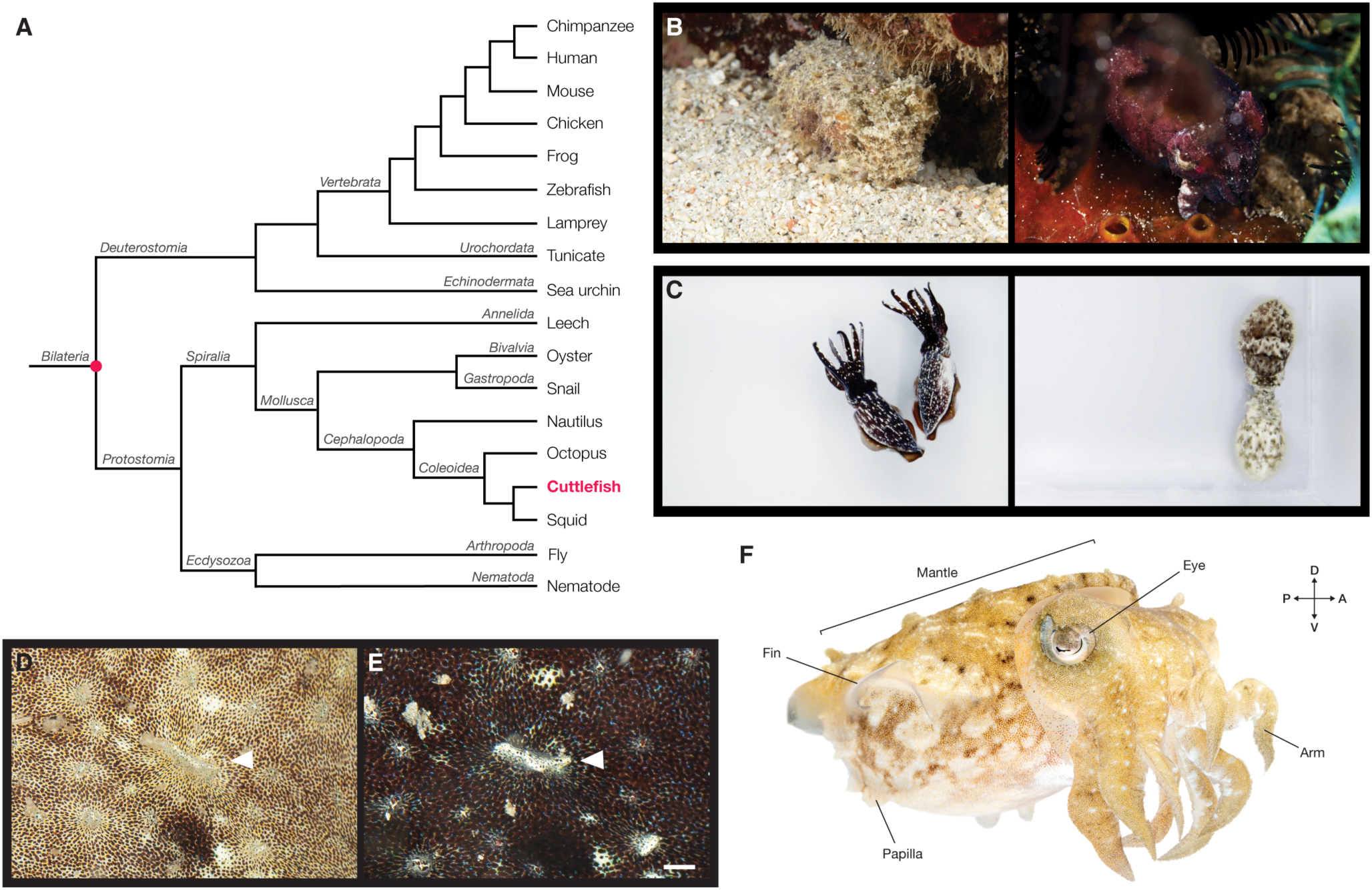
Introduction to the dwarf cuttlefish. A) Simplified phylogeny of Bilateria. Representative taxa, including widely used research organisms, are shown at branch tips to provide phylogenetic context. Cuttlefish belong to the coleoid cephalopod lineage of Mollusca. The pink circle marks the last common ancestor of cephalopods and vertebrates, which lived approximately 600 million years ago. Branch lengths are not to scale. B) Dwarf cuttlefish live in the Indo-Pacific and dynamically camouflage to their surroundings by adjusting the color, pattern, and texture of their skin. C) Dwarf cuttlefish use a repertoire of innate skin patterns in the presence of conspecifics that may reflect their internal state. Left: two males engage in a “dark aggression” skin pattern. Right: A male (top) and female (bottom) during mating. D,E) The same patch of cuttlefish skin before (D) and after (E) rapid color change. Chromatophores are flexible pigment sacs surrounded by radial muscles that are controlled by motor neurons projecting from the brain, enabling body-wide changes in color and pattern in less than a second. Papillae (arrowhead) can be rapidly extended or retracted to produce three-dimensional changes in skin texture and are also under direct neural control. Scale bar, 1 mm. F) Anatomy of a dwarf cuttlefish. The body (mantle) typically reaches 70–80 mm in length. Cuttlefish have eight arms, two feeding tentacles (hidden in this image), a pair of fins, and two eyes with a characteristic W-shaped pupil.

Recent advances in cephalopod husbandry^7–9^, genomics^10–14^, behavioral analysis^15, 16^, electrophysiology^17^, and gene knockouts^18, 19^ have made cephalopods increasingly tractable experimental systems. These tools have enabled mechanistic studies of the many biological innovations found in this lineage, including adaptive camouflage and social signaling, extensive RNA recoding^20–22^, remarkable regenerative capacity^23^, and chemotactile sensation^24–26^. Despite these advances, stable transgenesis has remained a major challenge, limiting the ability to visualize and manipulate cephalopod cells and neural circuits.

Several factors have impeded the development of transgenesis. First, robust multigenerational culture is difficult because most cephalopods are challenging to maintain in captivity, requiring specialized husbandry expertise. Second, transgenesis requires large numbers of fertilized eggs, yet most cephalopod species either have high fecundity and are difficult to maintain in captivity, or are amenable to multigenerational culture but produce relatively few offspring. Third, embryo manipulation is technically challenging. Many cuttlefish, squid, and bobtail embryos are enclosed within multiple layers of protective jelly (Figure S1A) and the blastodisc is small and thin relative to the yolk mass, making accurate, non-lethal targeting difficult^27^ (Figure S1B). Finally, the stable establishment of transgenesis requires both efficient genomic integration and knowledge of functional regulatory elements. Stable integration methods have not previously been reported in cephalopods and typically occur at low efficiency even in established model systems, necessitating large numbers of injected embryos. Moreover, functional promoters have not been identified in cephalopods, and the evolutionary divergence between cephalopods and traditional genetic model organisms suggests that many commonly used promoters may not function effectively in this lineage.

We have developed methods that enable stable transgenesis in the dwarf cuttlefish, *Ascarosepion bandense* (Figure 1F). We developed breeding and embryo manipulation protocols, generated genomic resources to identify candidate promoters and integration sites, and evaluated multiple methods for stable transgene integration. We successfully generated transgenic cuttlefish lines using both CRISPR-mediated and transposon-based approaches, with the transposon Minos producing the highest frequency of stable transgenesis. These F_1_ mScarlet transgenic embryos allowed the examination of embryonic development, and these methods provide a foundation for developing future transgenic lines expressing fluorescent proteins, reporters of neural activity, and optogenetic tools.

## Results

### Dwarf cuttlefish husbandry and embryology enable genetic manipulation

Establishing transgenesis in a new organism first requires understanding its reproductive biology and embryology. The dwarf cuttlefish (Figure 1F) is a small tropical cephalopod species^28^ that can be housed in mixed-sex groups and reaches sexual maturity in ∼4 months, among the shortest generation time of any cephalopod (Figure 2). During mating, cuttlefish embrace face-to-face, and the male transfers a mass of spermatophores into the buccal area of the female using a specialized arm called a hectocotylus^27^. The female can store sperm for hours to weeks until egg laying^29^, at which point ∼1–20 eggs are fertilized using stored sperm, wrapped in multiple layers of inky jelly, and attached to rock or coral to develop (Figure 2). Egg laying can occur at any time of day or night.

**Figure 2.**
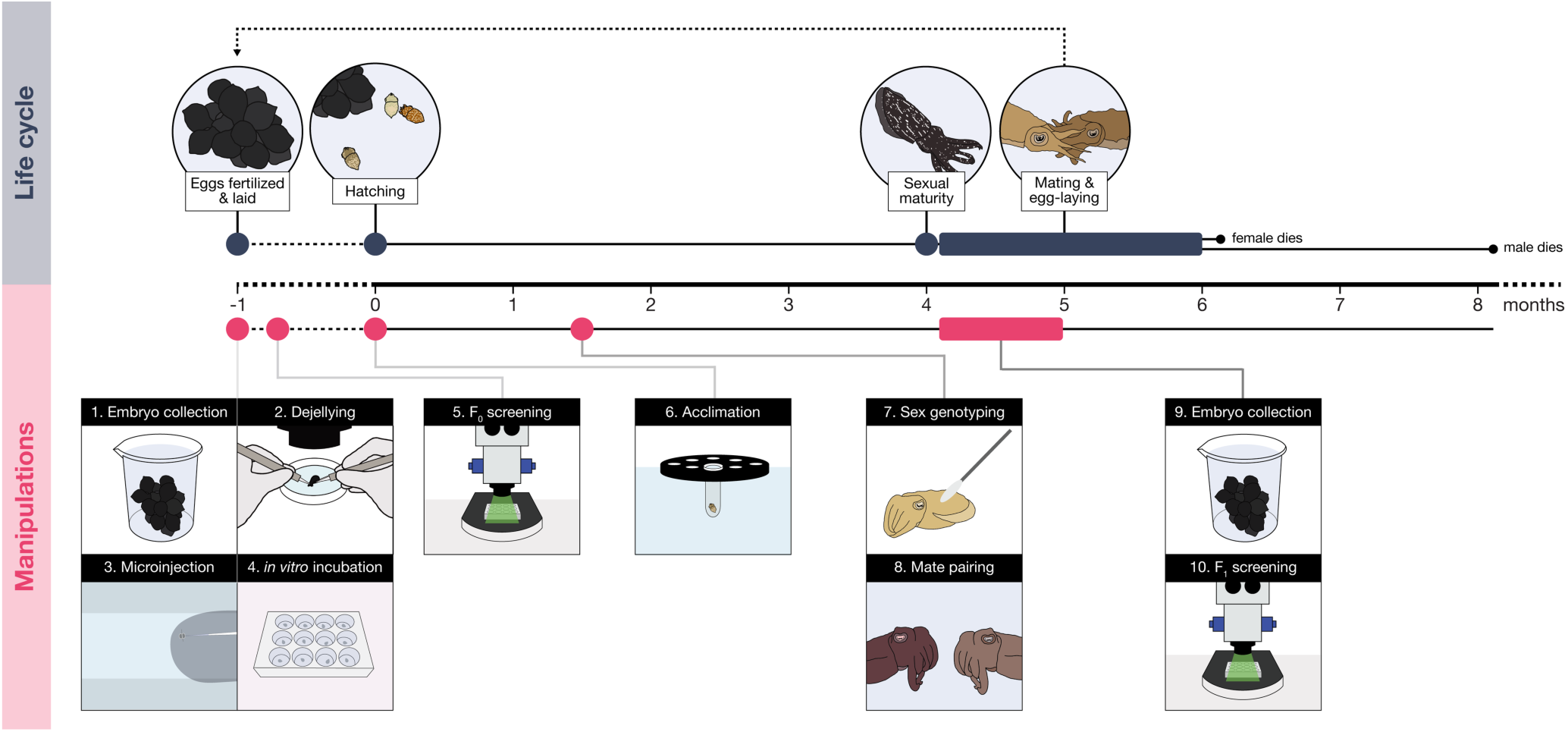
Dwarf cuttlefish husbandry and embryology enable genetic manipulation. **Life cycle:** Dwarf cuttlefish embryos develop within multiple layers of inky jelly and hatch after approximately one month. Hatchlings reach sexual maturity at ∼4 months and mate intermittently. Following mating, females store sperm for days to weeks before laying clutches of 1–20 eggs approximately every 5 days. Each egg is fertilized as it is laid, wrapped by the female in multiple layers of inky jelly, and attached to a rock or coral for development. Females typically live for ∼6 months, whereas males survive for ∼8 months. **Manipulations:** 1. To enable embryo microinjection, embryo collection is carried out every day. 2. Freshly laid embryos are sterilized in bleach and the many layers of jelly are manually removed using forceps. 3. Each embryo is injected in a custom made dish with beveled quartz needles. 4. Injected embryos are kept in agarose-coated 12-well dishes in an incubator with regular water changes until hatching (one month). 5. Embryos are screened for fluorescence at developmental stages 16-17 (8-9 days post-fertilization, dpf)^27^, when the body plan is established but the embryo remains relatively flat, allowing rapid assessment of transgene expression. 6. At hatching, each embryo is acclimated to the facility with gradual additions of system water. 7. At 6 weeks, the sex of each juvenile is determined using a skin swab and quantitative PCR^9,30^. 8. Sex genotyping enables mate pairing of each injected animal (F_0_) with a wild-type mate prior to sexual maturity. 9. After sexual maturity, at least six independent F_1_ clutches from each breeding group are screened for fluorescence to assess germline transmission. 10. Subsequent clutches from transmitting groups are reared with the surrounding jelly layers intact, while non-transmitting F_0_ animals are isolated.

Obtaining large numbers of one-cell-stage embryos for microinjection required optimizing dwarf cuttlefish breeding and egg collection. We established female-biased breeding groups using sex genotyping^9, 30^ and found that females preferentially laid eggs overnight, on weekends, and in visually sheltered locations. By exploiting these behavioral preferences through simple environmental manipulations (Methods), we shifted most egg laying into a predictable daytime window, enabling routine collection of one-cell-stage embryos (Figure 2). To enable microinjection of the thin blastodisc of the cuttlefish embryo, we adapted our embryo preparation protocol to remove the protective jelly^27^. Fertilization rates were similar between uninjected embryos maintained in the incubator after dejellying (104/116, 89.7%) and embryos maintained with their jelly intact in the animal facility (109/121, 90.1%). Among fertilized embryos, 58.7% (61/104) of dejellied embryos hatched, compared with 100.0% (109/109) of jelly-intact embryos. We developed an embryo holding dish and custom needles that enabled us to puncture the tough chorion (Figure 2; Figure S1C-E; Methods). The first embryonic cleavage occurs approximately 8 hours after fertilization^27^, providing a substantial window for microinjection of one-cell-stage embryos.

### CRISPR-mediated transgenesis

We initially pursued CRISPR-mediated transgenesis via targeted knock-ins. Our goal was to insert a P2A-transgene cassette immediately upstream of the stop codon of the targeted gene, allowing the transgene to be expressed from native regulatory elements. We generated transcriptomes from multiple adult tissues and initially selected broadly and highly expressed genes with well-characterized ubiquitous homologs. We reasoned that the expression of these genes would provide a visible readout of integration success in embryos and adults. Three candidates were selected: *actin I, actin II* (with homology to *beta-actin*), and *14-3-3 protein epsilon* (hereafter referred to as *epsilon*), a phosphoprotein-binding protein involved in diverse cellular processes, including cell signaling and cell-cycle regulation^31^ (Figure 3A; Tables S1, S2). Short guide RNAs (gRNAs) were designed to target sequences immediately upstream of the stop codon whenever permitted by the local genomic sequence (Methods; Table S1). Each gRNA was co-injected with recombinant Cas9 to assess editing efficiency. Embryos co-injected with the *actin I* gRNA developed abnormally and died during embryogenesis, potentially due to indel-induced frameshifts predicted to result in a 53-amino-acid C-terminal truncation (Methods; Table S1). Embryos co-injected with the *actin II* gRNA showed no detectable editing (Table S1), whereas embryos co-injected with the *epsilon* gRNA were edited efficiently without visible developmental defects (Table S1). Based on these results, *epsilon* was selected for knock-in experiments.

**Figure 3.**
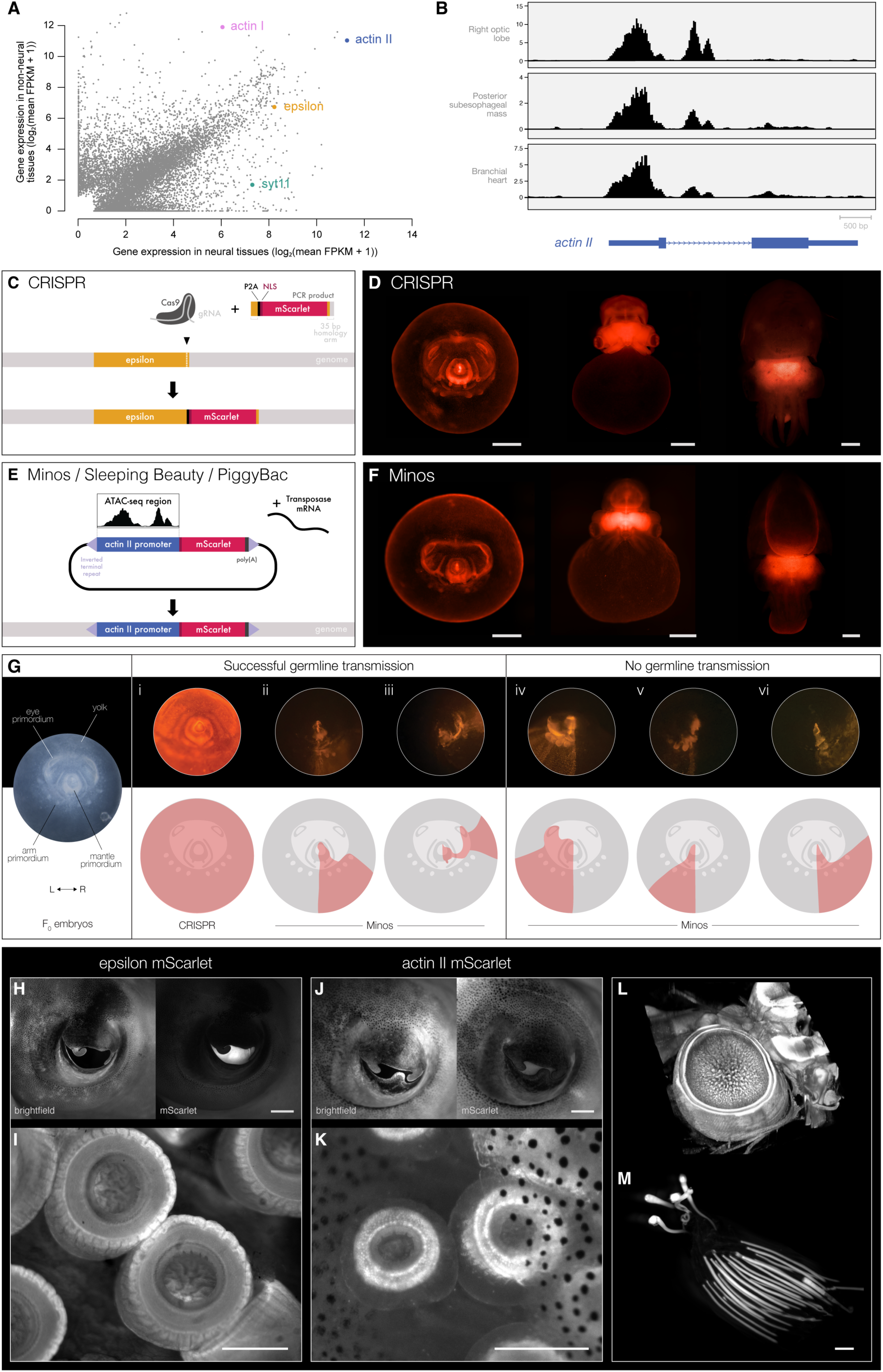
CRISPR- and transposon-mediated generation of stable transgenic cuttlefish. A) Scatter plot comparing gene expression in neural and non-neural adult dwarf cuttlefish tissues. Genes selected for CRISPR targeting or regulatory-element analysis (*actin I, actin II, epsilon*, and *syt11*) are highlighted. Values represent log₂(mean FPKM + 1) across neural or non-neural tissues. Gene sequences, gRNA targets, primers, and tissue-specific expression values are provided in Tables S1 and S2. B) ATAC-seq profiles at the *actin II* locus from branchial heart and two brain regions: right optic lobe and posterior subesophageal mass. Signal represents CPM-normalized chromatin accessibility. Y-axis limits were adjusted independently for each tissue to facilitate visualization of peak shape. Numeric axes show the corresponding CPM-normalized signal values. C) Strategy for CRISPR-mediated targeted knock-in. Cas9 and a guide RNA (gRNA) targeting the 3′ end of the *epsilon* coding sequence were co-injected with a PCR product encoding P2A-NLS-mScarlet flanked by 35-bp homology arms. The P2A peptide enables production of separate Epsilon and mScarlet proteins from a single transcript. NLS, nuclear localization sequence. D) F_1_ CRISPR *epsilon* mScarlet transgenic embryos at stages 17, 20, and 25 (hatching), left to right. Scale bars, 1 mm. E) Strategy for transposon-mediated transgenesis. The *actin II* regulatory element identified by ATAC-seq was cloned upstream of mScarlet between the inverted terminal repeats of the indicated transposon. The donor plasmid was co-injected with mRNA encoding the corresponding transposase. F) F_1_ Minos *actin II* mScarlet transgenic embryos at stages 17, 20, and 25 (hatching), left to right. Scale bars, 1 mm. G) All injected F_0_ embryos were imaged at stage 17 to record their fluorescence patterns. After sexual maturity, the F_1_ progeny of each animal were screened for fluorescence to determine whether the corresponding F_0_ animal transmitted the transgene through the germline. Stage 17 fluorescence images and schematic summaries are shown for representative F_0_ embryos that did (i–iii) or did not (iv–vi) exhibit germline transmission. H) Eye from an *epsilon* mScarlet adult, shown in brightfield (left) and mScarlet fluorescence (right). Scale bar, 1 mm. I) mScarlet fluorescence in suckers from an *epsilon* mScarlet adult. Scale bar, 500 μm. J) Eye from an *actin II* mScarlet adult, shown in brightfield (left) and mScarlet fluorescence (right). Scale bar, 1 mm. K) mScarlet fluorescence in suckers from an *actin II* mScarlet adult. Scale bar, 500 μm. L) Fixed and cleared brain from an *epsilon* mScarlet adult, imaged by light-sheet microscopy. M) mScarlet fluorescence in spermatophores from an *epsilon* mScarlet adult. Scale bar, 1 mm.

We adapted a short-homology donor strategy^32^ to generate a transgenic *epsilon* allele, using a PCR product encoding P2A-mScarlet^33^ with 35 bp homology arms flanking the gRNA target site to promote homology-directed repair (HDR) (Figure 3C; Methods). Co-injection of the donor with Cas9 and the gRNA produced mosaic fluorescent embryos in 11.3% of embryos (Table 1), which ranged from a few labeled cells to embryos exhibiting complete fluorescence. Sequencing of sub-regions of mosaic F_0_ embryos revealed that indels without transgene integration were present in cells in regions of the embryo that were not fluorescent (Figure S2). Survival of fluorescent injected embryos to hatching was extremely low (0.2%; Table 1); however, one F_0_ founder that exhibited complete fluorescence in the embryo, likely due to an insertion event at the one-cell-stage, survived to adulthood. Despite exhibiting morphological abnormalities – including reduced body size and an elongated cuttlebone – it successfully mated with a wild-type female, and transmitted the transgene through the germline, producing F_1_ fluorescent offspring (Figure 3D). Sequencing of the *epsilon* locus confirmed that the transgene was correctly integrated at the target locus, which deleted the four C-terminal amino acids of the Epsilon protein (Figure S3). F_1_ transgenic embryos exhibited widespread fluorescence throughout embryogenesis, with particularly bright signal in the brain (Figure 3D), and developed normally to adulthood. These experiments established the feasibility of stable transgenesis in cephalopods, albeit at extremely low efficiency (0.03% of injected embryos; Table 1). This revealed that poor embryo survival, rather than inefficient integration, was the principal limitation of the CRISPR approach (see Discussion). This motivated the evaluation of alternative transgenesis strategies.

**Table 1.** Comparison of CRISPR- and transposon-mediated transgenesis strategies in the dwarf cuttlefish. Numbers (n) and percentages (%) of embryos exhibiting fluorescence, surviving to hatching, surviving to sexual maturity, and transmitting the transgene through the germline. *Fluorescent embryos are reported as fluorescent / surviving embryos at the stage of fluorescence detection. All other percentages are relative to the total number injected.

|  | CRISPR |  | Sleeping Beauty |  | PiggyBac |  | Minos |  |
| --- | --- | --- | --- | --- | --- | --- | --- | --- |
|  | n | % | n | % | n | % | n | % |
| Embryos injected | 3429 |  | 330 |  | 343 |  | 236 |  |
| Fluorescent embryos* | 81 / 716 | 11.31% | 22 / 101 | 21.78% | 0 / 149 | 0.00% | 61 / 95 | 64.21% |
| Fluorescent survival to hatching | 7 | 0.20% | 11 | 3.33% | 0 | 0.00% | 22 | 9.32% |
| Fluorescent survival to mating | 1 | 0.03% | 10 | 3.03% | 0 | 0.00% | 14 | 5.93% |
| Transgenic germline | 1 | 0.03% | 3 | 0.91% | 0 | 0.00% | 3 | 1.27% |

### Transposon-mediated transgenesis

Transposons integrate at random genomic locations and require exogenous regulatory elements to drive transgene expression. Attempts to use heterologous promoters or the entire 4-kb region immediately upstream of the transcription start site of endogenous genes failed to drive detectable transgene expression (Methods). We therefore performed ATAC-seq to identify putative cuttlefish regulatory elements. At the *actin II* locus, ATAC-seq revealed a prominent accessibility peak (Figure 3B; Table S3) that overlapped the first exon and intron.

We first tested the *actin II* putative regulatory element in Tol2- and I-SceI-based reporter constructs. Tol2 produced no detectable fluorescence, whereas rare fluorescent cells were observed with I-SceI (Methods). We then evaluated Sleeping Beauty, Minos, and PiggyBac by placing the same reporter cassette between their inverted terminal repeats (Figure 3E). Each construct was co-injected at the one- or two-cell stage with mRNA encoding the corresponding transposase. In contrast to Tol2 and I-SceI, Sleeping Beauty and Minos produced robust mosaic reporter expression in 21.8% and 64.2% of embryos, respectively, whereas PiggyBac produced no detectable fluorescence (Table 1). Expression in Sleeping Beauty- and Minos-injected embryos ranged from small patches of labeled cells to embryos exhibiting complete fluorescence. The fluorescent embryos generated using Sleeping Beauty and Minos exhibited substantially higher survival to hatching (3.33% and 9.32%, respectively) than those generated by CRISPR-mediated knock-in (0.20%; Table 1). Importantly, germline transmission was successful, and occurred approximately 30-fold more frequently with Sleeping Beauty and 42-fold more frequently with Minos than with CRISPR (Table 1; Figure 3F). These results establish Sleeping Beauty and Minos as efficient methods for stable transgenesis in cuttlefish.

We next investigated whether early embryonic expression could predict germline transmission by prospectively imaging and tracking every mScarlet fluorescent embryo through adulthood. We observed large, continuous domains of fluorescence (Figure 3G), consistent with early transgene integration followed by the stereotyped contribution of individual blastomeres to the developing embryo. Notably, every founder that transmitted the transgene to the next generation exhibited fluorescence in the right posterior mantle primordium (Figure 3G), consistent with the approximate location of adult germ cells (Figure S4). One animal with right posterior mantle expression did not transmit the transgene, indicating that this pattern enriches for, but does not guarantee, germline transmission (Figure 3G-vi). Given the considerable effort and expense of raising all F_0_ cephalopods to adulthood, mating them, and screening their progeny, embryos with fluorescence in the right posterior mantle can therefore be prioritized in future transgenic line generation.

We examined mScarlet expression in adult F_1_ animals from the CRISPR *epsilon* and Minos *actin II* lines. Fluorescence was observed in the eyes, brain, spermatophores, and suckers (Figures 3H–M), with particularly bright expression in the eyes of *epsilon* mScarlet animals (Figure 3H). Notably, whereas *epsilon* mScarlet was broadly expressed throughout the sucker (Figure 3I), *actin II* mScarlet labeled a restricted population of cells in the suckers’ transition zone and infundibulum (Figure 3K), regions thought to contain chemotactile sensory cells^26,34^. These patterns demonstrate cell-specific transgene expression in adult tissues.

We next searched for neural-specific regulatory elements by searching for known neural genes with nearby ATAC-seq peaks that were enriched in neural relative to non-neural tissue (Methods). The *Synaptotagmin-11* (*syt11*) gene, encoding a synaptic protein^35^, revealed a prominent accessibility peak overlapping the first exon (Figure S5A; Table S3). We generated a dual-color Minos construct containing the *actin II* regulatory element driving superfolder GFP (sfGFP) and the *syt11* regulatory element driving mScarlet (Figure S5B). Co-injection of this construct with Minos transposase mRNA generated reporter expression in 38.0% of injected embryos (Figure S5C), despite the large 6.6 kb insert. Importantly, whereas GFP became detectable in the first few days of development and was often expressed across large regions of the body, mScarlet appeared later, and was restricted to the brain and other neural tissues only within GFP-positive regions (Figure S5C). We assessed expression of the dual-color construct in F_0_ embryos but did not test germline transmission. The tissue-restricted expression observed in these embryos demonstrates that ATAC-seq can be used to identify endogenous regulatory elements dictating neural-specific transgene expression in cuttlefish.

### Transgenic cuttlefish enable visualization of cellular dynamics *in vivo*

The nuclear-localized mScarlet of F_1_ transgenic embryos enabled live imaging of embryogenesis from the earliest stages of development. Under multiphoton microscopy, we captured the migration and fusion of the maternal and paternal pronuclei before the first cleavage (Figure 4A; Video 1). Unlike many vertebrate and invertebrate embryos, in which the maternal and paternal pronuclei migrate toward one another^36–39^, pronuclear fusion appeared to involve migration of a single pronucleus to the other (Figures 4A,C). After fusion, the fluorescence signal was lost, possibly due to nuclear envelope breakdown, after which the zygote underwent a series of stereotyped cleavage divisions that we reconstructed as a lineage (Figures 4B,C; Video 1). We also observed deviations from the process of pronuclear fusion. In one case, the first cleavage began before the pronuclei had fused. One pronucleus appeared to divide, while the lagging pronucleus changed trajectory and subsequently fused with one of the resulting nuclei (Figure S6; Video 2), indicating that the mechanism directing pronuclear fusion can persist beyond cell division. This embryo subsequently exhibited abnormal development.

**Figure 4.**
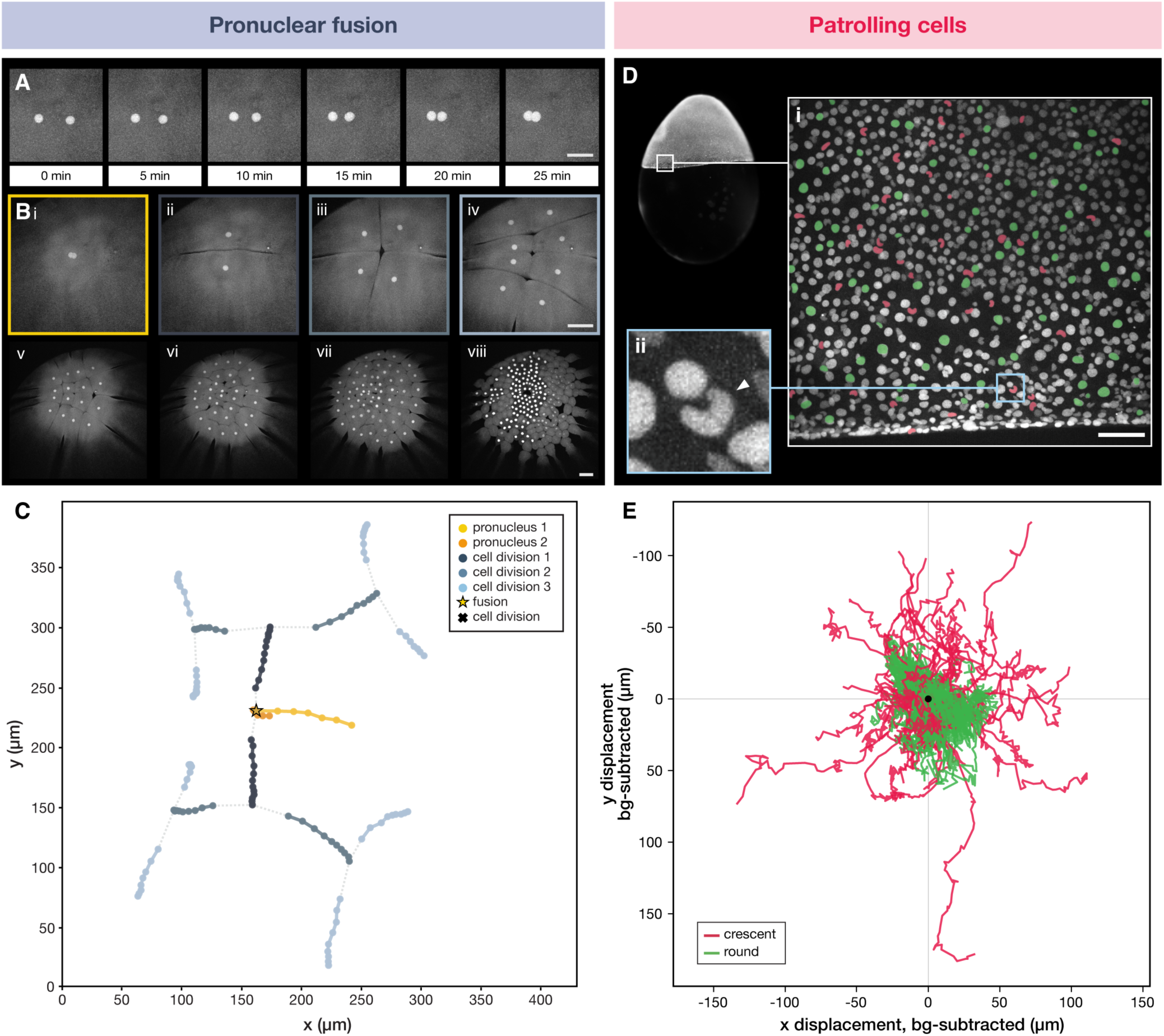
F_1_ transgenic cuttlefish enable in vivo visualization from embryogenesis through adulthood. A) Pronuclear fusion in a fertilized F_1_ transgenic embryo imaged by two-photon microscopy at 10-min intervals. Frames are from the time-lapse shown in Video 1 and analyzed in C. Time is shown relative to the 0-min frame, corresponding to the second acquired frame. Scale bar, 50 µm. B) Two-photon images from the same time-lapse showing (i) pronuclear fusion and (ii–viii) early cleavage of the mScarlet transgenic embryo. Early divisions occurred synchronously, with the onset of asynchronous division apparent in viii; nuclei are not visible in cells undergoing nuclear envelope breakdown. Panels i–iv correspond to the tracked lineages shown in C. Scale bars, 100 µm. See Video 1. C) Tracked lineages from the time-lapse shown in A and B (Video 1). The two pronuclei (yellow and orange) migrated together and fused (star), after which the zygote underwent successive cell divisions (division 1, dark blue; division 2, medium blue; division 3, light blue). D) Patrolling cells during epiboly. (i) First frame of the two-photon time-lapse, with tracked crescent-shaped nuclei false-colored pink (n = 61) and tracked round nuclei false-colored green (n = 118). Scale bar, 100 µm. For unlabeled image, see Figure S7A. See Video 3. (ii) Higher-magnification view showing a crescent-shaped nucleus (arrowhead) and round nuclei. E) Trajectories of cells with crescent-shaped nuclei (pink) and cells with round nuclei (green) after subtraction of embryo-wide movement.

We next used the transgenic line to visualize cellular dynamics during gastrulation. At three days post-fertilization, the blastoderm of the cuttlefish embryo begins spreading over the yolk in a process known as epiboly^27^. Most cells during epiboly contained round nuclei and moved coherently as a collective sheet (Figures 4D,E; Video 3). However, we identified a distinct population of cells with crescent-shaped nuclei that moved independently of this collective movement (Figures 4D,E; Figure S7; Video 3). Quantitative tracking revealed that cells with crescent-shaped nuclei moved more rapidly than cells with round nuclei and exhibited more directed movement, with superdiffusive motion (α = 1.39) compared with the approximately diffusive motion of cells with round nuclei (α = 0.96) (Figure S7). Although the identity of these migratory cells remains unknown, their distinctive nuclear morphology is reminiscent of cephalopod hemocytes^40^, and their behavior is reminiscent of recently described patrolling embryonic neutrophils in the annual killifish^41^, raising the possibility that they represent an early migratory, immune-like population. Together, these observations illustrate how stable transgenesis enables live imaging of previously inaccessible cellular behaviors during cuttlefish development.

## Discussion

We have established CRISPR and transposon-mediated strategies to generate stable transgenic cuttlefish. CRISPR-mediated knock-in strategies enabled precise modification of endogenous loci (Figures 3C,D) and should permit expression of transgenes at levels mirroring the targeted endogenous gene. However, we obtained only one transgenic cuttlefish after injection of over 3,000 embryos, a frequency 42-fold lower than we obtained with Minos (Table 1). We do not know whether these extremely low frequencies are unique to the *epsilon* gene or are a more general property of CRISPR-mediated transgenesis in cuttlefish. Targeting the *epsilon* gene generated mosaic fluorescent embryos at reasonable frequencies (11.3%), suggesting that the low frequency of embryo survival (0.20%), rather than initial integration efficiency, was the primary limitation of this approach. Uninjected dejellied embryos hatched at a rate of 58.7% (61/104), compared with 100% (109/109) of jelly-intact embryos maintained in the animal facility, indicating that *in vitro* embryo culture contributed to a baseline level of embryonic mortality.

Our strategy for targeting the *epsilon* gene was predicted to truncate the Epsilon protein by four C-terminal amino acids. It is possible, given the involvement of Epsilon in diverse cellular processes, this small C-terminal deletion could result in embryonic lethal phenotypes. Moreover, sequencing of spatially distinct regions in a separate CRISPR-targeting experiment (Figure S2) revealed indels without transgene integration in mosaic embryos, in regions lacking transgene expression, raising the possibility that widespread indels compromised development. However, the reduced survival during embryonic development may reflect a more general consequence of Cas9/gRNA injection rather than disruption of *epsilon* specifically: preliminary experiments targeting other genes not thought to be essential for development also showed extremely low survival following Cas9/gRNA injection (Table S4). Whatever the mechanism, the low frequency of survival following Cas9/gRNA injection, whether specific for *epsilon* or more generally for other genes in the cuttlefish, resulted in a prohibitively low yield of transgenic animals at the *epsilon* locus. We note, however, that CRISPR-induced mutagenesis was highly efficient, making this approach well suited for generating loss-of-function alleles when injection of the gRNA–Cas9 reagents does not cause lethality.

The transposons Sleeping Beauty and Minos yielded substantially higher embryo survival and germline transmission than CRISPR (Table 1). Furthermore, F_0_ adults routinely developed normally to adulthood, making Sleeping Beauty and Minos efficient vectors for routine transgenesis. Unlike CRISPR-mediated knock-in, however, the genomic locations of the integrated transgenes remain unknown. We have not analyzed the integration sites, and it is unknown whether mScarlet integrated at a single genomic locus in our founders, at multiple loci, or as tandem arrays. Notably, the transposon systems evaluated here originated from evolutionarily distant taxa: a salmonid fish (Sleeping Beauty)^42^, medaka fish (Tol2)^43^, moth (PiggyBac)^44^, and fly (Minos)^45^. Our results, together with prior work, demonstrate that transposase-based systems can function across broad evolutionary distances.

Successful transposon-mediated transgenesis required the identification of endogenous regulatory elements, for which ATAC-seq proved effective. Heterologous promoters and *A. bandense* upstream sequences failed to drive transgene expression (Methods). At the *actin II* locus, ATAC-seq identified a compact accessible region spanning the first exon and intron, outside the upstream sequence that had previously been tested. This ATAC-defined sequence (Figure 3B; Table S3) was sufficient to drive robust transgene expression without the addition of a known minimal promoter. Moreover, an ATAC-defined element from a neural-specific gene exhibited enhanced accessibility in neural tissue, and was sufficient to confer tissue-specific transgene expression (Figure S5; Table S3). The successful integration of the 6.6-kb dual-color construct further suggests that Minos can accommodate relatively large, multi-gene cargos in cuttlefish. Thus, ATAC-seq may provide a viable strategy for identifying cryptic regulatory elements, including those capable of tissue- or cell-specific transgene expression.

The establishment of stable transgenesis represents a major technological advance for cephalopod research. Together with the growing collection of publicly available resources for the dwarf cuttlefish^9, 27, 28, 30, 46, 47^, our transgenesis methods enable genetic approaches that were previously inaccessible, including the generation of lines with cell type-specific reporters, genetically encoded activity indicators, and optogenetic tools. The ability to stably alter endogenous genes and introduce novel genes into the genome of cephalopods should enable mechanistic investigation of the unique biology of these fascinating organisms.

## Methods

### Ethics statement

The use of cephalopods in laboratory research is currently not regulated in the USA. However, Columbia University has established strict policies for the ethical use of cephalopods, including operational oversight from the Institutional Animal Care and Use Committee (IACUC). All of the cuttlefish used in this study were handled according to an approved IACUC protocol (AC-AABT8673), including the use of deep anesthesia and the minimization and prevention of suffering.

### Animal husbandry

Dwarf cuttlefish (*Ascarosepion bandense*) were housed and bred at Columbia University in a recirculating artificial seawater system (30–35 parts per thousand (ppt); Red Sea Coral Pro Salt) maintained at 24–25°C. Water quality was maintained by regular water changes and mechanical, chemical, and biological filtration, with ammonia at 0 parts per million (ppm), nitrite <0.1 ppm, nitrate <30 ppm, and pH 8.0–8.4. Animals younger than 40 days post-hatching were fed live mysid shrimp (*Americamysis bahia*), after which they were transitioned to live grass shrimp (*Palaemonetes spp.*). Animals were fed three times daily.

Female-biased breeding groups consisting of four females and two males or five females and one male were established at six weeks post-hatching after sex determination by skin swab and quantitative PCR, following our published protocol (Figure 2)^9, 30^. Following sexual maturity (approximately 4 months post-hatching), cuttlefish mated intermittently, and each female laid fertilized eggs (embryos) every few days.

To increase the proportion of fertilized eggs laid during working hours, each breeding tank was set up with three artificial plants and one hollow resin tower (Imagitarium) that served as an egg-laying substrate^48^. At 9:30 am each day, a rock was placed over the tower to create a dark enclosure, which encouraged egg-laying. At 2:00 pm the rock was removed, deposited eggs were collected and transferred to plastic beakers containing artificial seawater, and the tower remained uncovered until 9:30 am the next day. Tanks were cleaned each afternoon by siphoning waste and debris from the bottom, which further delayed egg laying until the following morning.

### Embryo preparation for microinjection

After collection from the tanks, embryos with the surrounding jelly layers intact were surface sterilized by incubation in 0.5% sodium hypochlorite for 10 s, rinsed in 0.22-μm-filtered 31 ppt artificial seawater (FASW), incubated a second time in 0.5% sodium hypochlorite, neutralized in sodium thiosulfate (10 g/L), and rinsed twice in FASW with gentle swirling during the hypochlorite and thiosulfate incubations to ensure uniform exposure and facilitate chlorine removal. The surrounding jelly layers were manually removed with forceps in FASW in sterilized glass dishes. Dejellied embryos were transferred to agarose-coated 12-well plates containing 2 mm of 1% ultrapure agarose prepared in FASW and maintained in freshly prepared 1× antibiotic-antimycotic solution (A5955, Sigma-Aldrich) diluted in FASW. Embryos were maintained in 12-well plates at room temperature until they reached the one- or two-cell stage and were ready for injection.

### Microinjection needle fabrication

Injection needles were prepared from quartz glass capillaries containing an internal microfilament (1.00 mm outer diameter, 0.70 mm inner diameter, 10 cm length) using a Sutter P-2000 laser micropipette puller with the following settings: (Heat = 750; Fil = 4; Vel = 60; Del = 140; Pul = 175). Needles were beveled on a Sutter BV-10 beveler using a fine diamond abrasive plate (104D, Sutter) at a 15° angle for 25 s while maintaining positive pressure (using the Sutter Xenoworks digital injector) to prevent liquid from entering the needle during beveling. Needles were cleaned in 70% ethanol before storage.

### Embryo microinjection and incubation

Microinjections were performed using a Sutter Xenoworks digital injector, and Sutter MPC-200/MPC-325 micromanipulator. To immobilize embryos while maintaining positive pressure against the chorion during injection, a custom 3D-printed embryo holder (Extended Data 1) was placed in a 150-mm plastic dish and fitted with plastic semen straws (NC9780199, IMV International) that gently secured each embryo. Each embryo was injected with pressure 0; pulse width 0.5 s. Embryos were visualized during microinjection with darkfield illumination on a Zeiss SteREO Discovery V8 microscope.

Following injection, embryos were returned to agarose-coated 12-well plates and maintained at 28°C. No water changes were performed during the first ∼9 days of development (through stage 17)^27^. At stage 17, before chorion expansion, embryos were transferred with a glass pipette (14-955-501, Fisher Scientific) to agarose-coated 6-well plates. The pipette opening was first enlarged by removing the narrow tip with a ceramic tile (CTS, Sutter) and fire-polishing the edge with a Bunsen burner. From stage 17 onwards, 3 mL (20% volume) water changes were performed three times per week.

### RNA sequencing and transcriptome assembly

An adult dwarf cuttlefish (*Ascarosepion bandense*) was euthanized in 4% ethanol in artificial seawater (ASW). The vertical lobe, optic lobe, posterior subesophageal mass (PSM), stellate ganglion, retina, skin, sucker, and tentacle were rapidly dissected for RNA sequencing. Tissues were flash frozen in liquid nitrogen and stored at −80°C. RNA was extracted using TRIzol according to the manufacturer’s instructions. Total RNA was poly(A)-selected and sequenced on an Illumina HiSeq 2000.

Paired-end 100 bp RNA-seq reads from the eight tissues were aligned to the *A. bandense* reference genome (August 2021 assembly, https://www.cuttlebase.org/genome-transcriptome) using HISAT2^49^ v2.2.1. Aligned reads were converted to sorted BAM files using SAMtools^50^ v1.11. Because no reference transcript annotation was available, transcripts were assembled independently for each tissue using StringTie^51^ v2.1.4 with default parameters. Individual transcript assemblies were merged using StringTie (--merge) to generate a unified reference transcriptome, which was subsequently used for transcript quantification. Transcript abundances (FPKM) were estimated using StringTie and analyzed in R using Ballgown^51^ v2.40.0. Transcripts with low variance across tissues were removed using the criterion rowVars(texpr(bg)) > 1 before calculating gene-level expression. Mean expression was calculated across neural tissues (optic lobe, vertical lobe, PSM, stellate ganglion, and retina) and non-neural tissues (skin, sucker, and tentacle), and gene expression was plotted as log₂(mean FPKM + 1) in neural versus non-neural tissues. Genes were ranked by mean expression in neural tissues, and the exonic sequences of top-ranked candidates were extracted from the reference transcriptome and searched against known protein sequences using BLASTX (NCBI). *actin I*, *actin II*, and *14-3-3 protein epsilon* were selected as CRISPR targets based on their high expression, broad tissue distribution, and well-characterized homologs.

### ATAC-seq library preparation

Adult dwarf cuttlefish (*A. bandense*) were euthanized in 10% ethanol in artificial seawater (ASW), and branchial heart, optic lobes, and posterior subesophageal mass (PSM) were rapidly dissected for independent ATAC-seq experiments. Nuclei were isolated following the Omni-ATAC nuclei isolation protocol^52^. All steps were performed on ice, and all centrifugation steps were performed at 4°C. Briefly, fresh tissue was dounce homogenized in homogenization buffer (10 mM Tris pH 7.8, 0.1 mM EDTA, 5 mM CaCl₂, 3 mM Mg(OAc)₂, 320 mM sucrose, 0.1% NP-40, and 0.17 mM β-mercaptoethanol, supplemented with PMSF and protease inhibitors). Large tissue fragments were removed by centrifugation at 60 × g for 1 min. The remaining sample was layered onto an iodixanol density gradient and centrifuged at 3,000 × g for 20 min in a swinging-bucket rotor with the brake off. Nuclei free of visible cellular debris were collected from the interfacial band between the iodixanol layers and used for Omni-ATAC-seq transposition reactions.

For each tissue sample, 25,000–50,000 nuclei were resuspended in 50 µl transposition mixture (25 µl 2× TD buffer, 2.5 µl transposase [100 nM final], 16.5 µl PBS, 0.5 µl 1% digitonin, 0.5 µl 10% Tween-20, and 5 µl H₂O) and incubated at 37°C for 30 min with shaking at 1,000 rpm. Reactions were purified using the Zymo DNA Clean & Concentrator-5 Kit (Zymo Research). ATAC-seq sequencing adapters containing sample barcodes were added during five initial cycles of library amplification using NEBNext 2× Master Mix (New England Biolabs), and the number of additional amplification cycles required for each library was determined by qPCR. Libraries were purified using 1.6× volumes of AMPure XP beads and sequenced using 75-bp paired-end reads. Three biological replicates were performed for each tissue.

### ATAC-seq analysis

Paired-end ATAC-seq reads were aligned to the *A. bandense* genome using Bowtie2^53^ v2.4.4 with default settings, except that the maximum insert size was increased to 1,000 bp to allow alignment of larger fragments. Reads mapping to the mitochondrial genome were removed, and alignments were filtered to remove duplicate reads and retain reads with mapping quality ≥30, then converted to sorted, indexed BAM files. Filtered alignments were converted to tag directories and peaks were called using HOMER^54^ v4.11.1. For visualization, replicate accessibility profiles were visually inspected for concordance, after which BAM files from individual replicates were merged by tissue using SAMtools, and coverage tracks were generated using deepTools^55^ bamCoverage v3.5.6 with counts-per-million (CPM) normalization. Normalized tracks were visualized in the Integrated Genome Browser (IGB, v10.2.0). For comparison of peak shape across tissues, y-axis limits were adjusted independently while retaining the true CPM-normalized axis values.

Because CPM-normalized accessibility signal was lower in PSM and branchial heart than in the optic lobes, sequencing depth and library quality were examined as potential explanations. Peak counts normalized to total mapped reads confirmed that these differences persisted after depth normalization, and fragment-size distributions showed the expected nucleosome-associated periodicity in all tissues.

Differential chromatin accessibility was assessed by comparing left optic lobe and branchial heart ATAC-seq profiles using DiffBind^56^ v3.2.7. Peaks significantly enriched in the optic lobe relative to heart (FDR < 0.01, ≥8-fold change) were annotated to the nearest gene using HOMER annotatePeaks.pl. This gene list was cross-referenced with the RNA-seq expression data and ranked by neural tissue expression to identify highly expressed neural genes associated with neural-enriched accessible chromatin regions. Candidate genes were reviewed manually, and their exonic sequences were extracted from the reference transcriptome and searched against known protein sequences using BLASTX (NCBI) to confirm gene identity. *synaptotagmin-11* (*syt11*) was identified through this approach and selected as a candidate neural regulatory element.

### Regulatory element selection

Chromatin accessibility at each candidate locus was visualized in IGB using the merged, CPM-normalized accessibility tracks alongside the reference transcriptome annotation. Loci with a single prominent ATAC-seq peak associated with the candidate gene were prioritized to minimize ambiguity with neighboring regulatory elements. The genomic sequence underlying the selected ATAC-seq peak was used to design PCR primers for amplification from genomic DNA (see “Construction of transposon vectors”).

### CRISPR target design

The genomic sequences of *actin I, actin II,* and *epsilon* were identified in the cuttlefish genome and used to identify viable guide RNA (gRNA) targets in the 3’ end of the open reading frame (ORF), as close to the stop codon as possible, using CHOPCHOP^57–59^. gRNA targets were identified in *epsilon* with a predicted cutting site 11 bp away from the stop codon (GCAGGATGTGGGAACCGAGG), in *actin I* 163 bp from the stop codon (TCTTCATTGTGCTGGGGGCC) and in *actin II* 20 bp from the stop codon (GCACTTCCTATGGACGATAGAGG). gRNAs were synthesized by Synthego and co-injected with Cas9-NLS protein (M0646T, New England Biolabs) at the following concentrations: 1 μL of 5 mg/mL Cas9-NLS; 1 μL of 30 μM gRNA; 0.75 μL of water. To determine gRNA editing efficiencies, individual injected embryos were collected at stage 13 or later, the yolk was removed, and genomic DNA was extracted using the Monarch Spin gDNA Extraction Kit (T3010L, New England Biolabs). Genomic regions spanning each target site were amplified by PCR (150–500 bp amplicons) and submitted to Genewiz for Amplicon-EZ deep sequencing with SNP/indel detection analysis to quantify editing efficiency and identify the mutant alleles present in each embryo (Table S1).

### CRISPR-mediated integration

*mScarlet-I3*^33^ was amplified from a plasmid (gift from Aalok Varma and Loren Looger) using primers encoding 35-bp homology arms designed to direct in-frame integration at, or as close as possible to, the Cas9 cleavage site (3 bp upstream of the PAM): epsilon_mScarlet_F: AACAAAAAACTGAGCAGCTGCAGGATGTGGGAACCGGATCTGGTGCTACCAACTTCTC; epsilon_mScarlet_R: CAAACTAAATGGGCCACCGTTTATGACACCTCCTCTTAGCTGCCGCCGCTG. Primers were synthesized by IDT with a 5′ Amino Modifier C6 (5AmMC6), and the PCR product was purified using an E.Z.N.A. Cycle Pure Kit (Omega Bio-tek). Embryos were injected with a mix of 1 μL of 5 mg/mL Cas9-NLS, 1 μL of 30 μM gRNA, and 0.75 μL of 90 ng/μL PCR donor.

### Genotyping of CRISPR transgenics

Integration of mScarlet was initially identified by fluorescence. To confirm accurate integration, genomic DNA was isolated from either embryos (stage 13 or later following yolk removal) or skin swabs collected by gently stroking the dorsal mantle of juveniles or adults with a sterile foam swab. DNA was extracted using the Monarch Spin gDNA Extraction Kit (T3010L, New England Biolabs). Primers flanking the gRNA target site (epsilon_F: TGAATGTTAGTCTCACTGTATTAAAGTTAAAACC; epsilon_R: GTTCTCATATGAGTGTTGTTGTAATGTGC) amplified a 167-bp product from the wild-type allele and a 965-bp product following mScarlet integration. PCR products were separated on a 3% agarose gel, gel purified, and sequenced using the Plasmidsaurus PCR sequencing service to verify precise, in-frame integration at the target locus.

### Construction of transposon vectors

The 2.4 kb *actin II* regulatory sequence identified by ATAC-seq (Table S3) was synthesized by Eurofins Genomics. Because the sequence contains part of the first exon and open reading frame (ORF), the annotated *actin II* initiator ATG was mutated by site-directed mutagenesis to minimize translation from the *actin II* sequence. Three additional ATG sequences in the *actin II* ORF were also mutated to reduce the possibility of translation initiation at downstream sites and production of potentially toxic peptides. A P2A ribosome-skipping sequence was also included between the *actin II* sequence and NLS-mScarlet to separate any peptide initiated within the *actin II* sequence from the reporter. The modified *actin II* regulatory sequence was subsequently cloned upstream of P2A-NLS-mScarlet-I3-SV40p(A) between the ITRs of Sleeping Beauty, PiggyBac, and Minos reporter constructs, as described below. Neither reporter coding sequences nor transposase ORFs were codon optimized for *A. bandense*.

For Sleeping Beauty, ITR sequences were obtained from VectorBuilder (vector ID VB010000-9391ssw). P2A-NLS-mScarlet-I3-SV40p(A) and the right ITR were synthesized together by IDT. Because the inverted repeat structure prevented synthesis of both ITRs in a single construct, the left ITR was synthesized as two GeneBlocks and assembled into the plasmid by Gibson cloning. The 2.4 kb *actin II* regulatory sequence was subsequently inserted upstream of P2A by Gibson cloning.

For PiggyBac, left and right ITR sequences were provided by David Stern and synthesized by IDT flanking P2A-NLS-mScarlet-I3-SV40p(A). The 2.4 kb *actin II* regulatory sequence was subsequently inserted upstream of P2A by Gibson cloning.

For Minos, ITR sequences were provided by David Stern. P2A-NLS-mScarlet-I3-SV40p(A) and the right ITR were synthesized together by IDT. Because the inverted repeat structure prevented synthesis of both ITRs in a single construct, the left ITR was amplified from a GeneBlock and inserted by restriction digest and ligation. The 2.4 kb *actin II* regulatory sequence was subsequently inserted upstream of P2A by restriction digest and ligation.

To generate the dual-color Minos plasmid, the 1.2-kb *syt11* regulatory sequence identified by ATAC-seq (Table S3) was amplified from genomic DNA using primers syt11_F (GTAAATGATGTAGCAAGAGAAAACGTG) and syt11_R (GTGCGTGTATGTGTGTATTTGTG). Superfolder GFP (sfGFP)^60^ and a 100-bp zebrafish genomic sequence used as an insulator were synthesized by IDT. The *syt11* regulatory sequence driving mScarlet-I3 and the *actin II* regulatory sequence driving sfGFP were assembled in the Minos backbone, separated by the insulator sequence, using a combination of Gibson and conventional cloning.

### Transposase mRNA synthesis and transposon-mediated integration

The Sleeping Beauty SB100X transposase ORF was obtained from VectorBuilder (vector ID VB010000-9366jzq), while the PiggyBac and Minos transposase ORFs were provided by David Stern. Each transposase ORF was synthesized by IDT downstream of a *Xenopus laevis* β-globin 5′ UTR and upstream of an SV40 polyadenylation sequence in a pUC plasmid.

For *in vitro* transcription, each transposase plasmid was linearized with NotI and transcribed using the mMESSAGE mMACHINE SP6 Transcription Kit (Ambion). DNA templates were removed by DNase digestion, and RNA was purified using the E.Z.N.A. Total RNA Kit I (Omega Bio-tek).

For transposon-mediated integration, transposase mRNA and reporter plasmids were individually aliquoted (1 µL and 2 µL, respectively) and stored at −80°C and −20°C, respectively. Individual aliquots were thawed and combined immediately before injection. Embryos were injected with 1 µL of the corresponding Sleeping Beauty, PiggyBac, or Minos transposase mRNA at 100 ng/µL and 2 µL of the corresponding reporter construct at 50 ng/µL.

### Additional transgenesis approaches

As preliminary tests, Tol2 transposon- and I-SceI meganuclease-based approaches were also evaluated for transgenesis. Reporter constructs containing Tol2 ITRs or flanking I-SceI recognition sites were generated using a range of heterologous regulatory sequences, including *Drosophila hsp70*, human *HSP70*, mouse *ef1a*, human *EF1A*, human *ubiquitin*, *Crassostrea gigas* (oyster) *ef1a* (gift of Jonathan Henry), *Amphimedon queenslandica* (sponge) *isl2*, *tdrd3*, and *ccne1*, as well as approximately 4 kb of sequence upstream of the *actin I* or *actin II* start codon. None of these constructs produced detectable reporter fluorescence following embryo injection with Tol2 or I-SceI protein, and these approaches were not pursued further.

### F_0_ embryo fluorescence screening

Following injection with CRISPR or transposon reagents, embryos were initially screened for fluorescence between 3 and 5 days post-fertilization (dpf; stages 11–13)^27^ using a Zeiss SteREO Discovery V20 with Pentafluar microscope. For both *epsilon*- and *actin II*-driven mScarlet, fluorescence first became detectable at approximately 3 dpf. Fluorescent embryos were photographed at stage 17 (9 dpf) through the microscope eyepiece using an iPhone camera. At this stage, the body plan is established but the embryo remains relatively flat, allowing fluorescent cells to be visualized and their approximate positions within the body plan recorded before they become obscured by thicker tissues. Between stages 11 and 17, the numbers of fluorescent and surviving embryos were recorded to calculate the frequency of fluorescent embryos. Fluorescent embryos that survived to hatching were acclimated to the facility (see “Embryo acclimation to the facility”).

### Embryo acclimation to the facility

At hatching (stage 25)^27^, when the majority of the yolk had been consumed and only a small remnant remained visible between the arms, each hatchling was transferred from the culture dish to a 2-mL microcentrifuge tube filled halfway with water (∼1 mL). Tubes were transported to the animal facility, placed in a foam tube rack, and floated in a rearing tank. After 10 min to equilibrate to the tank temperature, 100 μL of tank water was added to each tube and incubated for an additional 10 min. This step was repeated until the tube was completely filled with tank water (∼1 hour). Once acclimation was complete, the tube was placed on its side in the tank, allowing the hatchling to swim out voluntarily. Hatchlings that had not emerged within 3 h were gently released by carefully tilting the tube. For embryos exhibiting developmental abnormalities, the timing of acclimation was adjusted. For instance, those with delayed development were maintained in the culture dish, and those that prematurely lost their yolk were acclimated to the facility the following day.

Hatchlings were not fed during the first two days post-hatching. Beginning on day 3, each animal was offered one live mysid shrimp (*Americamysis bahia*) daily. Food quantity was gradually increased every 10 days until animals were transitioned to live grass shrimp (*Palaemonetes spp.*) at 40 days post-hatching.

### F_0_ animal mate pairing and breeding

At six weeks post-hatching, the sex of injected animals was determined by skin swab and quantitative PCR following our published protocol (Figure 2)^9, 30^. Fluorescent F_0_ animals were then paired with wild-type mates. Where possible, mates were selected to be similar in size and to have hatched within 3–4 days of one another. F_0_ males were paired with one or two wild-type females, whereas F_0_ females were paired with a single wild-type male. All pairs were established by introducing the male into the tank occupied by the female. F_0_ animals exhibiting morphological abnormalities were paired with younger, smaller wild-type mates to reduce competition and aggression.

Following sexual maturity and the onset of egg laying, embryos were collected from each breeding group, dejellied, and maintained in 12-well plates in the incubator until 3 days post-fertilization (dpf), when they were screened for fluorescence. At least six independent clutches from each breeding group were screened to assess germline transmission of the transgene. For breeding groups exhibiting germline transmission, subsequent clutches were collected with the surrounding jelly layers intact and reared in the animal facility. Embryos were individually separated from the egg cluster and suspended in custom-made cradles constructed from ¼-inch plastic mesh or 3D-printed components within a recirculating aquatic system. Water flow was directed over the embryos to maintain gentle movement within the cradles. Once per week, embryos were dipped in 0.5% sodium hypochlorite for 3 s, neutralized in sodium thiosulfate (10 g/L), rinsed twice in artificial seawater (ASW), and returned to their cradles. This treatment was discontinued at 21 dpf to prevent premature hatching. Undeveloped embryos were removed as necessary to minimize biofouling.

### Embryo fluorescence imaging

A 1.2% solution of ultrapure low-melting-point agarose (Thermo Fisher Scientific, #16520100) in 0.22-μm-filtered artificial seawater (FASW) was maintained at 40°C, poured into a shallow Petri dish, and allowed to cool to room temperature. Once the agarose began to gel, the chorions were removed from embryos at stages 17, 20, and 25^27^ using forceps, and the embryos were transferred into the agarose using a glass transfer pipette. Embryos were imaged with a Zeiss SteREO Discovery V20 with Pentafluar microscope and Zeiss Axiocam 506 Mono camera. Fluorescence images were false-colored using a custom Fiji^61^ v2.16.0 macro. Brightness and contrast were adjusted uniformly across images in Adobe Photoshop. For stage 25 embryos, gamma was adjusted to visualize both the body and brain.

### Embryo time-lapse imaging

Embryos were embedded in 1.2% ultrapure low-melting-point agarose (ThermoFisher #16520100) dissolved in artificial seawater (35 ppt artificial reef salt in reverse osmosis water). The mix was kept at 40°C then poured into a shallow petri dish and allowed to cool to room temperature. Once the agarose began to gel, an embryo was added by a glass transfer pipette and positioned with the animal pole upwards to image pronuclear fusion and early cell division, or rotated 90 degrees (lateral view) to image epiboly. The dish was fixed with modeling clay to the bottom of a larger glass well containing artificial seawater.

Embryos were imaged on a Bergamo II multiphoton microscope with a Coherent Chameleon laser tuned to 1050 nm and a Nikon 16×/0.80 NA water-immersion objective (CFI75 LWD 16X W). The glass dish containing the embryo was placed on a slide warmer (Amscope TCS-200) previously set to maintain the water temperature at approximately 28°C. Once positioned, the gaps between the objective and the edges of the glass well were covered with parafilm to minimize evaporation. Embryos were imaged with z-stacking plus time-lapse at 5-min intervals, for a maximum of 8 hours. Recordings were visualized and z-projected in ImageJ and analyzed via custom Python scripts.

### Image analysis

Two-photon time-lapse stacks were maximum-intensity projected along the z-axis at each time point. For the epiboly motility analysis, nuclei were selected, tracked, and classified using custom Python tools (NumPy and Matplotlib). To obtain an unbiased spatial sample, the blastoderm was divided into a 10 × 10 grid, and one nucleus was selected from each occupied grid cell in the first frame. Selected nuclei were then tracked frame-by-frame by manually annotating their centroids. These manually generated trajectories were supplemented with automatically generated trajectories that were subsequently proofread. For automated tracking, nuclear centroids were detected using Otsu thresholding followed by intensity- and distance-weighted watershed segmentation and linked across frames by nearest-neighbor tracking after compensating for the coherent movement of the cell sheet. Nuclear morphology was manually classified for each tracked cell as crescent-shaped, round, or ambiguous.

To isolate cell-autonomous movement from the bulk movement associated with epiboly, the mean displacement vector of all tracked cells at each frame was subtracted from the displacement of each individual cell. Mean cell speed was calculated as the total path length divided by the total tracking time; cells tracked for fewer than 20 frames were excluded. Mean squared displacement (MSD) was calculated as the ensemble time-averaged squared displacement as a function of frame lag, and the anomalous-diffusion exponent α was estimated as the slope of log(MSD) versus log(lag). Displacements and speeds were converted to physical units using the acquisition pixel size and frame interval. Mean-speed distributions of cells with crescent-shaped and round nuclei were compared using a one-sided Mann–Whitney U test.

Pronuclear migration and cleavage divisions were tracked using an analogous automated cell-tracking tool. Nuclei were segmented using white-top-hat filtering, and linked across frames based on the overlap between nuclear masks across frames. Pronuclear fusion was identified by the overlap of two nuclear masks. After cell division, reformed nuclei were identified as the descendants of the nearest parent nuclear track.

### Adult live imaging

Adult F_1_ transgenic animals were anesthetized in a mixture of 175 g/L MgCl_2_ and 2% ethanol for 60 s and then placed in a glass dish under a Zeiss SteREO Discovery V20 with Pentafluar microscope. Images were captured using Zen software and brightness and contrast were adjusted in ImageJ. Spermatophores were dissected from a geriatric male cuttlefish after euthanasia in a mixture of 175 g/L MgCl_2_ and 10% ethanol.

### Tissue clearing and light-sheet imaging

Dissected brains were fixed for 24 h in 4% paraformaldehyde in artificial seawater, rinsed for 1 h in deionized water, incubated for 72 h in 50% tetrahydrofuran in deionized water, rinsed again for 1 h in deionized water, and incubated for 72 h in high-refractive-index EZ View solution, with gentle agitation throughout, following the EZ Clear tissue-clearing protocol^62^. For imaging, samples were transferred to a quartz cuvette containing custom refractive-index-matched Cargille immersion oil (RI 1.5120) and imaged on a Miltenyi Biotec UltraMicroscope II light-sheet microscope with support from the Zuckerman Institute’s Cellular Imaging Platform and the National Institutes of Health (NIH 1S10OD023587-01). Light-sheet data were visualized using Dragonfly 3D World image processing and analysis software.

## Data availability

RNA-seq and ATAC-seq deposited at X. Plasmids X deposited in Addgene.

## Author contributions

T.G.M. conceived the project, developed microinjection methods, performed the molecular cloning, CRISPR and transposon experiments, embryo microinjections, genotyping, F_0_ screening and characterization, germline-transmission screening, and characterization of stable transgenic lines, and analyzed the RNA-seq, ATAC-seq, and transgenesis data. C.J.G. developed the husbandry methods with input from T.J.M., K.A.P., and T.G.M. C.J.G., T.J.M., and K.A.P. maintained the cuttlefish colony and performed animal husbandry and transgenic breeding. G.T.B. performed the F1 embryo time-lapse imaging, adult live imaging, light-sheet imaging of cleared brains, and associated image analyses. E.V.B. acquired the ATAC-seq data and provided initial ATAC-seq analysis scripts. Y.X. and E.C.C. prepared embryos for injection and, together with T.G.M. and C.J.G., contributed to embryology troubleshooting. A.N. conceived the embryo holding mechanism. A.V. designed and introduced point mutations into the *actin II* ATAC-seq regulatory element in a plasmid generated by T.G.M. C.B.A. acquired the RNA-seq data. T.G.M., L.L.L., S.L., and R.A. provided supervision. T.G.M. wrote the original draft of the manuscript and created the figures with input from G.T.B. T.G.M. and R.A. reviewed and edited the manuscript with input from all authors. R.A. acquired funding for the project.

## Acknowledgements

We are grateful to Martin Mueller and Tanya Tabachnik of the Zuckerman Institute Advanced Instrumentation platform for designing and fabricating the embryo holder; members of the Axel lab, David Stern, Alexandre Paix, Andrés Bendesky, Josh Rosenthal, Philip Abitua, Fred Rubino, Nipam Patel, Phillip Cleves, David Hain, and Simone Rencken for helpful discussions; David Stern, Gary Struhl, and Jonathan Henry for sharing reagents; the Zuckerman Institute Cellular Imaging platform for instrument use and technical support; and Josh Rosenthal and the Marine Biological Laboratory’s Cephalopod Program for supporting the generation of the RNA-seq dataset for this project. This project was funded by a Howard Hughes Medical Institute Hanna H. Gray Fellowship (T.G.M.), an NSF Graduate Research Fellowship (G.T.B.), and the Howard Hughes Medical Institute (R.A.).

## Use of AI

During the preparation of this work, the authors used ChatGPT for editing suggestions to improve the clarity and legibility of the manuscript and Claude code to generate custom Python scripts for data annotation, analysis, and visualization. After using these tools, the authors reviewed and edited the content as needed and take full responsibility for the content of the published article.

## References

1. dos Reis, M., Thawornwattana, Y., Angelis, K., Telford, M.J., Donoghue, P.C., and Yang, Z. (2015). Uncertainty in the Timing of Origin of Animals and the Limits of Precision in Molecular Timescales. Curr Biol, 25, 2939–2950. 10.1016/j.cub.2015.09.066.

2. Jozet-Alves, C., Bertin, M., and Clayton, N.S. (2013). Evidence of episodic-like memory in cuttlefish. Curr Biol, 23, R1033–5. 10.1016/j.cub.2013.10.021.

3. Fiorito, G., von Planta, C., and Scotto, P. (1990). Problem solving ability of Octopus vulgaris Lamarck (Mollusca, Cephalopoda). Behav Neural Biol, 53, 217–230. 10.1016/0163-1047(90)90441-8.

4. Montague, T.G. (2023). Neural control of cephalopod camouflage. Curr Biol, 33, R1095–R1100. 10.1016/j.cub.2023.08.095.

5. Shook, E.N., Barlow, G.T., Garcia-Rosales, D., Gibbons, C.J., and Montague, T.G. (2024). Dynamic skin behaviors in cephalopods. Curr Opin Neurobiol, 86, 102876. 10.1016/j.conb.2024.102876.

6. Messenger, J.B. (2001). Cephalopod chromatophores: neurobiology and natural history. Biol Rev Camb Philos Soc, 76, 473–528. 10.1017/s1464793101005772.

7. Jolly, J., Hasegawa, Y., Sugimoto, C., Zhang, L., Kawaura, R., Sanchez, G., Gavriouchkina, D., Marlétaz, F., and Rokhsar, D. (2022). Lifecycle, culture, and maintenance of the emerging cephalopod models Euprymna berryi and Euprymna morsei. Frontiers in Marine Science, 9, 1039775. 10.3389/fmars.2022.1039775.

8. Grearson, A.G., Dugan, A., Sakmar, T., Sivitilli, D.M., Gire, D.H., Caldwell, R.L., Niell, C.M., Dölen, G., Wang, Z.Y., and Grasse, B. (2021). The lesser pacific striped octopus, Octopus chierchiae: an emerging laboratory model. Frontiers in Marine Science, 8, 1771. https://www.frontiersin.org/articles/10.3389/fmars.2021.753483/full?utm_source.

9. Rubino, F.A., Coffing, G.C., Gibbons, C.J., Small, S.T., Desvignes, T., Pessutti, J., Petersen, A.M., Arkhipkin, A., Shcherbich, Z., Postlethwait, J.H., et al. (2026). A non-invasive method to genotype cephalopod sex by quantitative PCR. iScience, 29, 115594. 10.1016/j.isci.2026.115594.

10. Coffing, G.C., Tittes, S., Small, S.T., Songco-Casey, J.O., Piscopo, D.M., Pungor, J.R., Miller, A.C., Niell, C.M., and Kern, A.D. (2025). Cephalopod sex determination and its ancient evolutionary origin. Curr Biol, S0960-9822(25)00005. 10.1016/j.cub.2025.01.005.

11. Gavriouchkina, D., Tan, Y., Parey, E., Ziadi-Künzli, F., Hasegawa, Y., Piovani, L., Zhang, L., Sugimoto, C., Luscombe, N., Marlétaz, F., et al. (2025). A single-cell atlas of the bobtail squid visual and nervous system highlights molecular principles of convergent evolution. Nat Ecol Evol, 10.1038/s41559-025-02720-9.

12. Albertin, C.B., Medina-Ruiz, S., Mitros, T., Schmidbaur, H., Sanchez, G., Wang, Z.Y., Grimwood, J., Rosenthal, J.J.C., Ragsdale, C.W., Simakov, O., et al. (2022). Genome and transcriptome mechanisms driving cephalopod evolution. Nat Commun, 13, 2427. 10.1038/s41467-022-29748-w.

13. Schmidbaur, H., Kawaguchi, A., Clarence, T., Fu, X., Hoang, O.P., Zimmermann, B., Ritschard, E.A., Weissenbacher, A., Foster, J.S., Nyholm, S.V., et al. (2022). Emergence of novel cephalopod gene regulation and expression through large-scale genome reorganization. Nat Commun, 13, 2172. 10.1038/s41467-022-29694-7.

14. Rencken, S.D., Tushev, G., Hain, D., Ciirdaeva, E., Simakov, O., and Laurent, G. (2026). Chromosome-scale genome assembly of the European common cuttlefish Sepia officinalis. Elife, 14, RP107393. 10.7554/eLife.107393.

15. Renard, M.D.M., Ukrow, J., Elmaleh, M., Evans, D.A., Wu, Y., Liang, X., and Laurent, G. (2026). Disentangling cephalopod chromatophores motor units with computer vision. Elife, 15, RP110074. 10.7554/eLife.110074.

16. Ukrow, J., Renard, M.D.M., Moghimi, M., and Laurent, G. (2025). A computational pipeline to track chromatophores and analyze their dynamics. Elife, 14, RP106509. 10.7554/eLife.106509.

17. Mano, T., Tsaridis, K., Kojima, Y., Masucci, G.D., Dinh, T.T.V., Tong, R., Glykos, V., Dolezalova, L., Asada, K., Shumkova, D., et al. (2026). Hierarchical processing and polarization encoding in the cephalopod visual system. bioRxiv, 2026.01. 08.698312. 10.64898/2026.01.08.698312.abstract.

18. Crawford, K., Diaz Quiroz, J.F., Koenig, K.M., Ahuja, N., Albertin, C.B., and Rosenthal, J.J.C. (2020). Highly Efficient Knockout of a Squid Pigmentation Gene. Curr Biol, 30, 3484–3490.e4. 10.1016/j.cub.2020.06.099.

19. Ahuja, N., Hwaun, E., Pungor, J.R., Rafiq, R., Nemes, S., Sakmar, T., Vogt, M.A., Grasse, B., Diaz Quiroz, J., Montague, T.G., et al. (2023). Creation of an albino squid line by CRISPR-Cas9 and its application for in vivo functional imaging of neural activity. Curr Biol, S0960-9822(23)00739. 10.1016/j.cub.2023.05.066.

20. Birk, M.A., Liscovitch-Brauer, N., Dominguez, M.J., McNeme, S., Yue, Y., Hoff, J.D., Twersky, I., Verhey, K.J., Sutton, R.B., Eisenberg, E., et al. (2023). Temperature-dependent RNA editing in octopus extensively recodes the neural proteome. Cell, 186, 2544–2555.e13. 10.1016/j.cell.2023.05.004.

21. Rangan, K.J., and Reck-Peterson, S.L. (2023). RNA recoding in cephalopods tailors microtubule motor protein function. Cell, 186, 2531–2543.e11. 10.1016/j.cell.2023.04.032.

22. Rosenthal, J.J.C., and Eisenberg, E. (2023). Extensive Recoding of the Neural Proteome in Cephalopods by RNA Editing. Annu Rev Anim Biosci, 11, 57–75. 10.1146/annurev-animal-060322-114534.

23. Imperadore, P., and Fiorito, G. (2018). Cephalopod tissue regeneration: consolidating over a century of knowledge. Frontiers in physiology, 9, 593. https://www.frontiersin.org/articles/10.3389/fphys.2018.00593/full.

24. Villar, P.S., Jiang, H., Shugaeva, T., Berdan, E.L., Kulkarni, A., Hiroi, M., Masucci, G., Reiter, S., Lindahl, E., Howard, R.J., et al. (2026). A sensory system for mating in octopus. Science, 392, 96–101. 10.1126/science.aec9652.

25. Sepela, R.J., Jiang, H., Shin, Y.H., Hautala, T.L., Clardy, J., Hibbs, R.E., and Bellono, N.W. (2025). Environmental microbiomes drive chemotactile sensation in octopus. Cell, 188, 4849–4860.e21. 10.1016/j.cell.2025.05.033.

26. van Giesen, L., Kilian, P.B., Allard, C.A.H., and Bellono, N.W. (2020). Molecular Basis of Chemotactile Sensation in Octopus. Cell, 183, 594–604.e14. 10.1016/j.cell.2020.09.008.

27. Montague, T.G., Rieth, I.J., and Axel, R. (2021). Embryonic development of the camouflaging dwarf cuttlefish, Sepia bandensis. Dev Dyn, 250, 1688–1703. 10.1002/dvdy.375.

28. Gibbons, C.J., Rubino, F.A., Barlow, G.T., Garcia-Rosales, D., Guevara, N., Rosales, B., Aneja, S., Elkis, D., Mendoza, G.D., Demayo, J.I., et al. (2025). Natural Habitat and Wild Behaviors of the Dwarf Cuttlefish, Ascarosepion bandense. Ecol Evol, 15, e72001. 10.1002/ece3.72001.

29. Hanlon, R.T., Ament, S.A., and Gabr, H. (1999). Behavioral aspects of sperm competition in cuttlefish, Sepia officinalis (Sepioidea: Cephalopoda). Marine Biology, 134, 719–728. 10.1007/s002270050588.

30. Rubino, F.A., Gibbons, C.J., Mungioli, T.J., Small, S.T., Coffing, G.C., Kern, A.D., and Montague, T.G. (2026). Protocol for genotyping cephalopod sex using a skin swab and quantitative PCR. STAR Protoc, 7, 104693. 10.1016/j.xpro.2026.104693.

31. Zhang, Y., Yan, M., Yu, Y., Wang, J., Jiao, Y., Zheng, M., and Zhang, S. (2024). 14-3-3ε: a protein with complex physiology function but promising therapeutic potential in cancer. Cell Commun Signal, 22, 72. 10.1186/s12964-023-01420-w.

32. Seleit, A., Aulehla, A., and Paix, A. (2021). Endogenous protein tagging in medaka using a simplified CRISPR/Cas9 knock-in approach. Elife, 10, e75050. 10.7554/eLife.75050.

33. Bindels, D.S., Haarbosch, L., van Weeren, L., Postma, M., Wiese, K.E., Mastop, M., Aumonier, S., Gotthard, G., Royant, A., Hink, M.A., et al. (2017). mScarlet: a bright monomeric red fluorescent protein for cellular imaging. Nat Methods, 14, 53–56. 10.1038/nmeth.4074.

34. Graziadei, P.P., and Gagne, H.T. (1976). Sensory innervation in the rim of the octopus sucker. J Morphol, 150, 639–679. 10.1002/jmor.1051500304.

35. Shimojo, M., Madara, J., Pankow, S., Liu, X., Yates, J., Südhof, T.C., and Maximov, A. (2019). Synaptotagmin-11 mediates a vesicle trafficking pathway that is essential for development and synaptic plasticity. Genes Dev, 33, 365–376. 10.1101/gad.320077.118.

36. Schatten, G. (1981). The movements and fusion of the pronuclei at fertilization of the sea urchin Lytechinus variegatus: Time-lapse video microscopy. J Morphol, 167, 231–247. 10.1002/jmor.1051670207.

37. Reinsch, S., and Karsenti, E. (1997). Movement of nuclei along microtubules in Xenopus egg extracts. Curr Biol, 7, 211–214. 10.1016/s0960-9822(97)70092-7.

38. Strome, S., and Wood, W.B. (1983). Generation of asymmetry and segregation of germ-line granules in early C. elegans embryos. Cell, 35, 15–25. https://www.cell.com/cell/fulltext/0092-8674(83)90203-9.

39. Scheffler, K., Uraji, J., Jentoft, I., Cavazza, T., Mönnich, E., Mogessie, B., and Schuh, M. (2021). Two mechanisms drive pronuclear migration in mouse zygotes. Nat Commun, 12, 841. 10.1038/s41467-021-21020-x.

40. Castellanos-Martínez, S., Prado-Alvarez, M., Lobo-da-Cunha, A., Azevedo, C., and Gestal, C. (2014). Morphologic, cytometric and functional characterization of the common octopus (Octopus vulgaris) hemocytes. Dev Comp Immunol, 44, 50–58. 10.1016/j.dci.2013.11.013.

41. Sattler, S.M., Lammers, N.C., Hriscu, S.V., Booth, C.L.T., Trapnell, C., and Abitua, P.B. (2026). Early immune cell development precedes gastrulation in annual killifish. bioRxiv, 2026.06. 25.734555. 10.64898/2026.06.25.734555.abstract.

42. Ivics, Z., Hackett, P.B., Plasterk, R.H., and Izsvák, Z. (1997). Molecular reconstruction of Sleeping Beauty, a Tc1-like transposon from fish, and its transposition in human cells. Cell, 91, 501–510. https://www.cell.com/fulltext/S0092-8674(00)80436-5.

43. Koga, A., Suzuki, M., Inagaki, H., Bessho, Y., and Hori, H. (1996). Transposable element in fish. Nature, 383, 30. 10.1038/383030a0.

44. Cary, L.C., Goebel, M., Corsaro, B.G., Wang, H.G., Rosen, E., and Fraser, M.J. (1989). Transposon mutagenesis of baculoviruses: analysis of Trichoplusia ni transposon IFP2 insertions within the FP-locus of nuclear polyhedrosis viruses. Virology, 172, 156–169. 10.1016/0042-6822(89)90117-7.

45. Franz, G., and Savakis, C. (1991). Minos, a new transposable element from Drosophila hydei, is a member of the Tc1-like family of transposons. Nucleic acids research, 19, 6646. https://pmc.ncbi.nlm.nih.gov/articles/PMC329244.

46. Lorig-Roach, R., Meredith, M., Monlong, J., Jain, M., Olsen, H.E., McNulty, B., Porubsky, D., Montague, T.G., Lucas, J.K., Condon, C., et al. (2024). Phased nanopore assembly with Shasta and modular graph phasing with GFAse. Genome Res, 34, 454–468. 10.1101/gr.278268.123.

47. Montague, T.G., Rieth, I.J., Gjerswold-Selleck, S., Garcia-Rosales, D., Aneja, S., Elkis, D., Zhu, N., Kentis, S., Rubino, F.A., Nemes, A., et al. (2023). A brain atlas for the camouflaging dwarf cuttlefish, Sepia bandensis. Curr Biol, 33, 2794–2801.e3. 10.1016/j.cub.2023.06.007.

48. Villanueva, R., Darmaillacq, A.-S., Kuba, M.J., Roumbedakis, K., Nabhitabhata, J., Almansa, E., Caamal-Monsreal, C., Court, M., Dickel, L., and Durante, E.D. (2026). Collection, handling and care of cephalopod eggs. Reviews in Fish Biology and Fisheries, 36, 17. 10.1007/s11160-025-10017-0.

49. Kim, D., Paggi, J.M., Park, C., Bennett, C., and Salzberg, S.L. (2019). Graph-based genome alignment and genotyping with HISAT2 and HISAT-genotype. Nat Biotechnol, 37, 907–915. 10.1038/s41587-019-0201-4.

50. Danecek, P., Bonfield, J.K., Liddle, J., Marshall, J., Ohan, V., Pollard, M.O., Whitwham, A., Keane, T., McCarthy, S.A., Davies, R.M., et al. (2021). Twelve years of SAMtools and BCFtools. Gigascience, 10, giab008. 10.1093/gigascience/giab008.

51. Pertea, M., Kim, D., Pertea, G.M., Leek, J.T., and Salzberg, S.L. (2016). Transcript-level expression analysis of RNA-seq experiments with HISAT, StringTie and Ballgown. Nat Protoc, 11, 1650–1667. 10.1038/nprot.2016.095.

52. Corces, M.R., Trevino, A.E., Hamilton, E.G., Greenside, P.G., Sinnott-Armstrong, N.A., Vesuna, S., Satpathy, A.T., Rubin, A.J., Montine, K.S., Wu, B., et al. (2017). An improved ATAC-seq protocol reduces background and enables interrogation of frozen tissues. Nat Methods, 14, 959–962. 10.1038/nmeth.4396.

53. Langmead, B., and Salzberg, S.L. (2012). Fast gapped-read alignment with Bowtie 2. Nat Methods, 9, 357–359. 10.1038/nmeth.1923.

54. Heinz, S., Benner, C., Spann, N., Bertolino, E., Lin, Y.C., Laslo, P., Cheng, J.X., Murre, C., Singh, H., and Glass, C.K. (2010). Simple combinations of lineage-determining transcription factors prime cis-regulatory elements required for macrophage and B cell identities. Mol Cell, 38, 576–589. 10.1016/j.molcel.2010.05.004.

55. Ramírez, F., Ryan, D.P., Grüning, B., Bhardwaj, V., Kilpert, F., Richter, A.S., Heyne, S., Dündar, F., and Manke, T. (2016). deepTools2: a next generation web server for deep-sequencing data analysis. Nucleic Acids Res, 44, W160–5. 10.1093/nar/gkw257.

56. Stark, R., and Brown, G. (2011). DiffBind: differential binding analysis of ChIP-Seq peak data. Bioconductor, 10.18129/B9.bioc.DiffBind.

57. Montague, T.G., Cruz, J.M., Gagnon, J.A., Church, G.M., and Valen, E. (2014). CHOPCHOP: a CRISPR/Cas9 and TALEN web tool for genome editing. Nucleic Acids Res, 42, W401–7. 10.1093/nar/gku410.

58. Labun, K., Montague, T.G., Gagnon, J.A., Thyme, S.B., and Valen, E. (2016). CHOPCHOP v2: a web tool for the next generation of CRISPR genome engineering. Nucleic Acids Res, 44, W272–6. 10.1093/nar/gkw398.

59. Labun, K., Montague, T.G., Krause, M., Torres Cleuren, Y.N., Tjeldnes, H., and Valen, E. (2019). CHOPCHOP v3: expanding the CRISPR web toolbox beyond genome editing. Nucleic Acids Res, 47, W171–W174. 10.1093/nar/gkz365.

60. Pédelacq, J.D., Cabantous, S., Tran, T., Terwilliger, T.C., and Waldo, G.S. (2006). Engineering and characterization of a superfolder green fluorescent protein. Nat Biotechnol, 24, 79–88. 10.1038/nbt1172.

61. Schindelin, J., Arganda-Carreras, I., Frise, E., Kaynig, V., Longair, M., Pietzsch, T., Preibisch, S., Rueden, C., Saalfeld, S., Schmid, B., et al. (2012). Fiji: an open-source platform for biological-image analysis. Nat Methods, 9, 676–682. 10.1038/nmeth.2019.

62. Hsu, C.-W., Cerda, J., Kirk, J.M., Turner, W.D., Rasmussen, T.L., Flores Suarez, C.P., Dickinson, M.E., and Wythe, J.D. (2022). EZ Clear for simple, rapid, and robust mouse whole organ clearing. Elife, 11, e77419. 10.7554/eLife.77419.

