## Supplementary material for "Generation of a transgenic cephalopod": Figure S

### Supplementary Materials

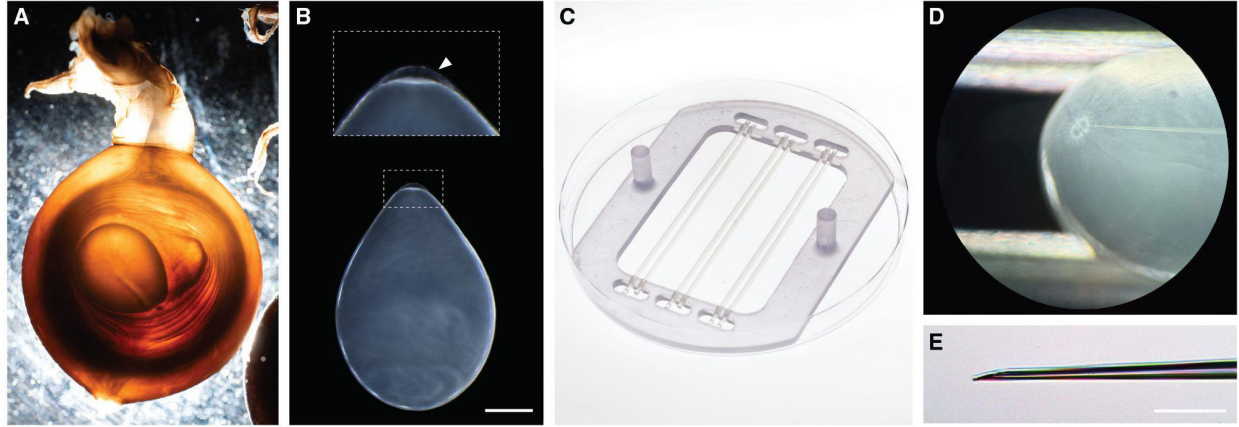

**Figure S1. Cuttlefish embryos and microinjection**

A) Dwarf cuttlefish embryos are enclosed within multiple layers of pigmented jelly. Removal of the outer opaque layer reveals the underlying layers seen here.

B) Cuttlefish embryo at the one-cell stage (darkfield image adapted from<sup>1</sup>). The small, flattened cell (inset) sits at the animal pole of the large yolk. The embryo is surrounded by a transparent chorion (arrowhead), visible here immediately above the cell. Scale bar, 1 mm.

C) Custom injection dish. Cuttlefish embryos are immobilized between bovine semen straws secured within a 3D-printed dish (Extended data 1).

D) Two-cell-stage embryo during microinjection. Because the chorion thickens at the animal pole, the needle is introduced at an angle to facilitate chorion penetration.

E) Quartz injection needles are beveled to 15° to penetrate the chorion. Scale bar, 100 μm.

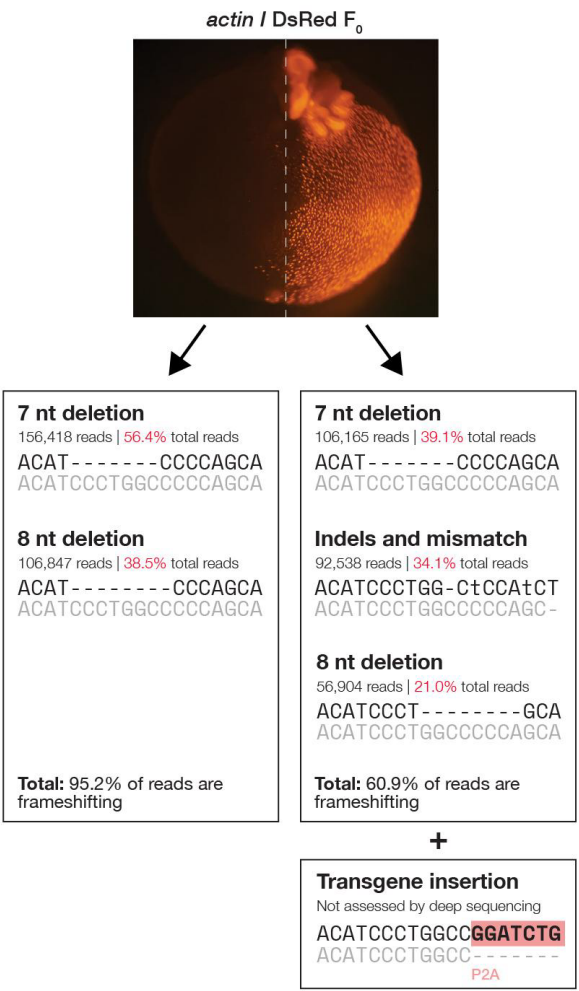

**Figure S2. Widespread indel formation in a mosaic CRISPR-targeted embryo.**

An embryo injected with Cas9, a gRNA targeting *actin* *I*, and a donor PCR product encoding P2A-DsRed exhibited fluorescence in approximately half of its body. To assess editing in the absence of transgene integration, fluorescent and non-fluorescent regions were dissected separately, and the endogenous *actin* *I* locus was amplified and deep sequenced. Indels were abundant in both regions, including the non-fluorescent region, indicating that widespread editing can occur in cells without transgene integration. Note that correct transgene integration was not assessed by deep sequencing because the primers flanked the insertion site and successful integration would have increased the amplicon by ~1 kb – beyond the size range used for amplicon sequencing, with PCR also expected to preferentially amplify the shorter unintegrated allele.

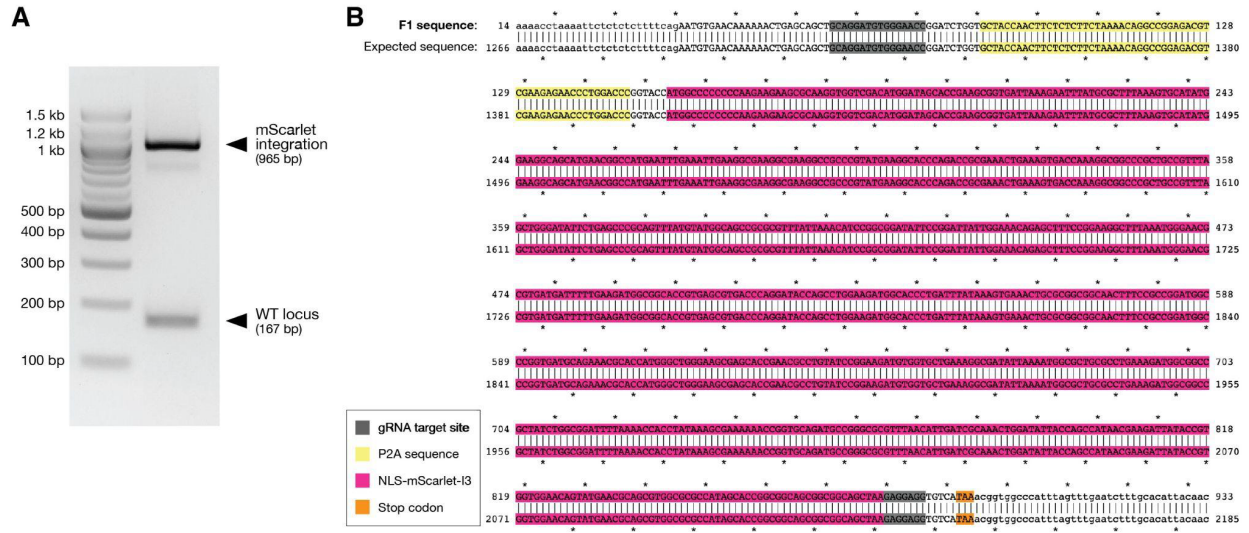

Figure S3. Confirmation of transgene integration at the epsilon locus.

A) The region surrounding the gRNA target site in epsilon was amplified from genomic DNA of a CRISPR F1 transgenic animal. As expected for a heterozygous animal, two products were generated: a 167-bp product corresponding to the unedited locus inherited from the wild-type parent and a 965-bp product corresponding to the mScarlet-containing allele.

B) Sequencing of the 965-bp product confirmed insertion of P2A (yellow) and NLS-mScarlet (pink) within the gRNA target site (dark gray), 12 nt upstream of the endogenous epsilon stop codon (orange), resulting in a four-amino-acid C-terminal truncation of Epsilon.

Figure S4. Anatomical location of the gonads in adult dwarf cuttlefish.

A) Wild-type female. The ovary, containing developing eggs, is located at the posterior end of the animal.

B) epsilon mScarlet transgenic male. The testis is located at the posterior end of the animal. The pink coloration likely reflects high mScarlet expression.

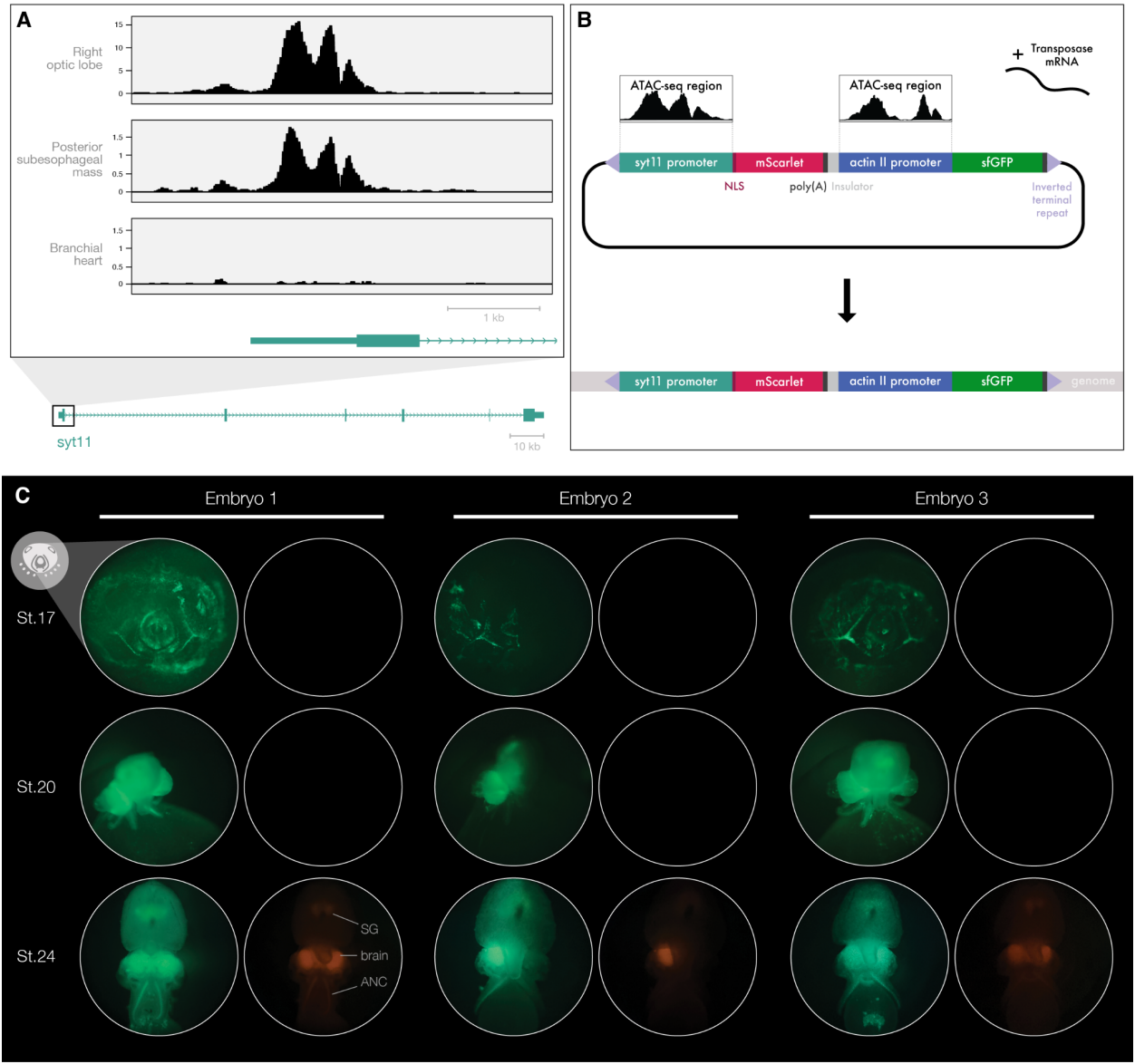

**Figure S5. An ATAC-seq-defined element at the *synaptotagmin-11* (*sytt11*) locus drives neural-specific expression.**

A) ATAC-seq profiles at the *sytt11* locus from branchial heart and two brain regions: right optic lobe and posterior subesophageal mass. Because the *sytt11* locus spans 142 kb, ATAC-seq profiles are shown at higher magnification around the first exon. Signal represents CPM-normalized chromatin accessibility. Y-axis limits were adjusted independently for each tissue to facilitate visualization of peak shape. Numeric axes show the corresponding CPM-normalized signal values.

B) Strategy for dual-color Minos transposon-mediated transgenesis. The *sytt11* regulatory element identified by ATAC-seq was cloned upstream of mScarlet, and the *actin II* regulatory element identified by ATAC-seq was cloned upstream of superfolder GFP (sfGFP). The two expression cassettes were separated by an insulator and cloned between the Minos inverted terminal repeats. The donor plasmid was co-injected with Minos transposase mRNA. NLS, nuclear localization sequence.

C) A total of 177 embryos were injected with the dual-color Minos construct. Of the 55 embryos surviving to the time of initial fluorescence screening, 21 (38.0%) exhibited GFP expression. Three representative embryos are shown. Top row: stage 17 (~9 days post-fertilization, dpf); middle row: stage 20 (~15 dpf); bottom row: stage 24 (~3.5 weeks post-fertilization, wpf). GFP expression was detectable at stages 17 and 20, whereas mScarlet was not detectable.

until stage 24, when it was restricted to the brain and other neural structures within GFP-positive regions. Neural structures are labeled in the stage 24 image of embryo 1: brain, stellate ganglion (SG), and axial nerve cord (ANC).

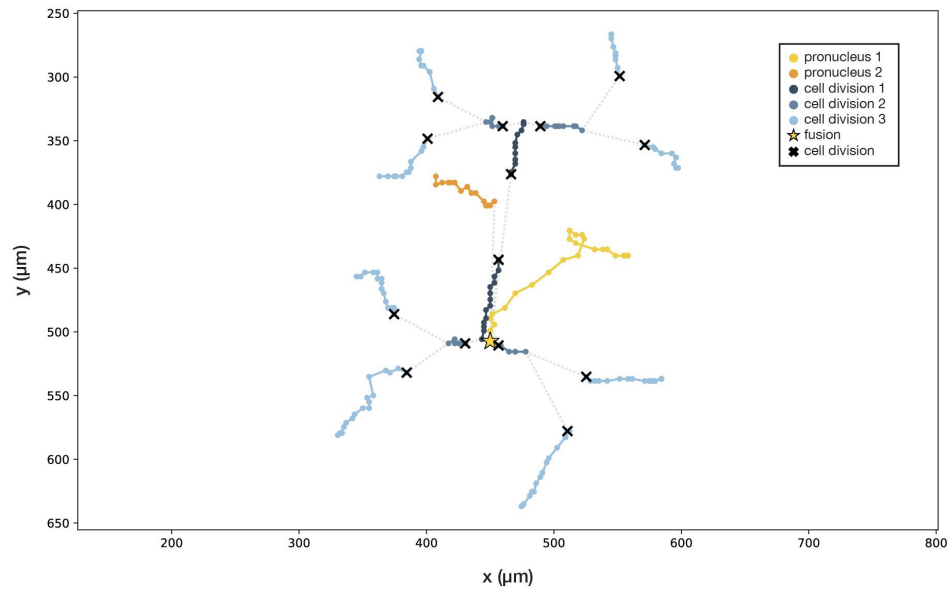

**Figure S6. Delayed pronuclear fusion.**

Manually tracked lineages of the two pronuclei (yellow and orange). The yellow pronucleus did not reach the orange pronucleus before the first cell division (dark blue), but subsequently changed its trajectory and fused with it (star) before the second cell division (medium blue). Division 3, light blue. Crosses mark division events. The embryo arrested development after a small number of divisions. See Video 2.

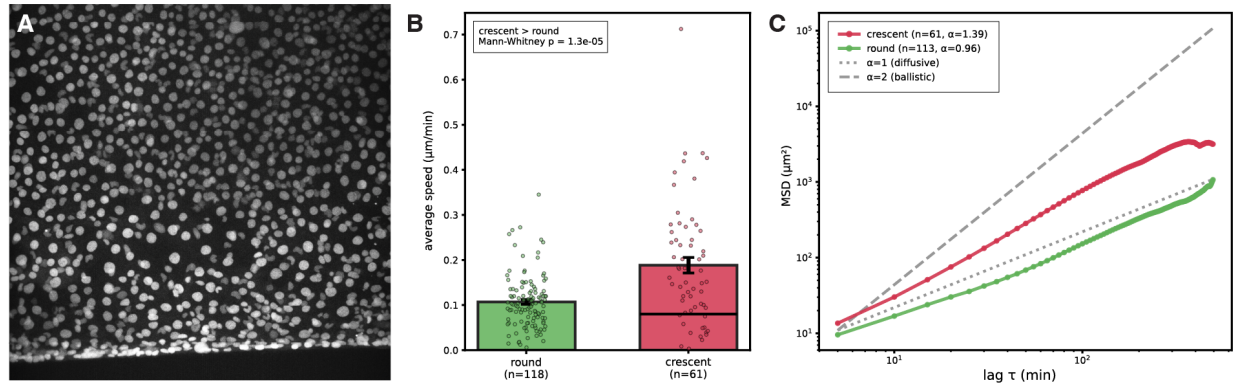

**Figure S7. Quantification of cell migration during epiboly according to nuclear morphology.**

A) Unmodified frame from the two-photon time lapse of cells during epiboly. For pseudocoloring of tracked round and crescent-shaped nuclei, see Figure 4Di.

B) Mean cell speed according to nuclear morphology. Cells with crescent-shaped nuclei ($n = 61$) moved significantly faster than cells with round nuclei ($n = 118$) (one-sided Mann–Whitney U test, $p = 1.3 \times 10^{-5}$).

C) Mean squared displacement (MSD) of cells with crescent-shaped and round nuclei. Scaling exponents ($\alpha$) are shown for each population, with $\alpha = 1$ indicating diffusive motion and $\alpha > 1$ indicating superdiffusive motion. Dashed reference lines indicate $\alpha = 1$ (diffusive) and $\alpha = 2$ (ballistic) motion.

For B and C, $n = 61$ cells with crescent-shaped nuclei and 113 cells with round nuclei were tracked for $\geq 20$ frames. Images were acquired at $0.407 \mu\text{m}/\text{pixel}$ every 5 min.

| Gene | Genomic location (Aug 2021 assembly on Cuttlebase) | Open reading frame (gRNA target in blue) | Amino acid sequence | gRNA target (5'→3') | Primers for sequencing | Cut rate | Dominant alleles |
| --- | --- | --- | --- | --- | --- | --- | --- |
| *actin I* / MSTRG.17104 | scaffold778:2708836-2711805 | ATGGTCGGTATGGGTCAGAAAGACTCCTACGTAGGAGACGAGGCCCA GAGCAAGAGAGGTATTCTTACCTTGAAATACCCCATTGAGCACGGTAT TGTCACCAACTGGGATGACATGGAGAAAATCTGGCATCACACATTCTA CAACGAGCTCCGTGTCGCTCCTGAAGAGCACCCCGTCCTTCTCACAG AGGCCCCCTTGAACCCCAAGGCTAACAGGGAAAAGATGACTCAAATCA TGTTCGAGACCTTCAACGCTCCCGCCATGTATGTTGCCATCCAGGCTG TCCTCTCTCTGTACGCTTCTGGTGTGTACAACCGGATTGTTCTTGATT CCGGAGACGGTGTACCCCACTGTACCAATCTATGAAGGTTATGCCCT TCCCCACGCCATCCTCCGTCTTGACTTGGCCGACGTGATCTTACTGA CTACCTCATGAAGATCCTTACCAGACGTGGTTATTCATTACAACCAACC GCCGAGAGAGAAATTGTGCGTGACATCAAGGAGAAATTGCTATGTC GCTCTTGACTTTGAACAGGAGATGGCCACTGCTGCTTCTTCTTCTCC CTCTCGAAGAGCTACGAGTTGCCCGATGGTCAAGTAATCACCATTGGT AACGAGAGGTTCAAGTGCCCGAATCTTGTTCACGCTTCTCTTCTTG GGTATGGAATCTGCTGGTATCCATGAGACAACATACAACCTCCATCATG AAATGCGATGTCGACATCAGGAAAGATCTGTATGCTAACACTGTGCTG TCTGGAGGTACCACCATGTTCCCTGGTATTGCTGACAGGATGCAGAAG GAAATCACATC**CCTGGCCCCCAGCACAATGAAGA**TCAAGATCATCGC TCCCCCCGAGCGTAAATACTCCGTCTGGATTGGCGGTTCCATCTTGGC TTCTCTGTCCACCTTCCAACAGATGTGGATTAGCAAGCAGGAATATGA CGAGTCCGGTCCATCTATCGTTCACCGCAAATGTTTCTAA | MVGMGQKDSYVGDEAQSKRGILTL KYPIEHGIVTNWDDMEKIWHHTFYN ELRVAPEEHPVLLTEAPLNPKANRE KMTQIMFETFNAPAMYVAIQAVLSLY ASGRRTTGIIVDSGDVHTTVPYIEG YALPHAILRLDLAGRDLTDYLMKILTE RGYSFTTTAEREIVRDIKEKLCYVAL DFEQEMATAASSSSLEKSYELPDGQ VITIGNERFRCPESLFQPSFLGMESA GIHETTYNSIMKCDVDIRKDLYANTV LSGGTTMFPGIADRMQKEITSLAPST MKIKIAPPERRYKYSVWIGGSILASLST FQQMWISKQEYDESGPSIVHRKCF* | TCTTCATTGTGCTGGGGCC**AGG** | **actin-I_F:**<br>ATCAGGAAAGATCTG TATGCTAACACTG | 96.99% mutant reads | **7 nt deletion:** CCCTGGC<br>**Result:** 53 amino acid truncation |
|  |  |  |  |  | **actin-I_R:**<br>TCATATTCCTGCTTG CTAATCCACATC |  | **8 nt deletion:** CCCTGGCC<br>**Result:** 43 amino acid truncation |
| *actin II* / MSTRG.58 | scaffold100:3586777-3591070 | ATGGATGATGAAGTTGCTGCTCTCGTTATTGACAACGGATCTGGTATG TCGAAAGCTGGTTTTGCCGGTGATGATGCACCTAGGGCTGATTCCCA TCTATTGTCGGCCGTCGCCGACATCAGGGTGTGATGGTGGTATGGG ACAGAAAGACAGATTATGTTGGTGATGAAGCTCAGAGCAAGAGAGGTAT TCTTACCTTGAAGTACCCTATTGAACACGGTATTGTCACAAAATGGGAT GACATGGAGAAAAATTTGGCATCATCTTCTATAATGAATTGAGAATTG CTCCTGAAGAGCATCCAGTTCTGTTAACAGAGCTCCTCTTAATCCTAA GGCGAACAGAGAAAAAGATGACTCAATCATGTTTGAACCTTCAACAC CCCTGCTATGTATGTGGCCATCCAAGCTGTCTTGCCCTATATGCTTCC GGTCTGACCACTGGTATTGTCATGGATTCTGGTGATGGTGCACTCAC ACTGTCCCATCTACGAAGGTTACGCCCTTCTCTATGCTATCCTCCGT TTGGATTGGCCGGCCGTGACTTGACAGATTACCTTATGAAGATCTTG ACCGAACGTGGTTACTCTTTCAACAACCACTGCTGAGAGAGAAATTGTC CGTGACATCAAGGAAAACTTTGCTATGTGGCTCTAGACTTCGATCAG GAAATGCAAAACAGCTGCTTCTTCTCTTCAATTGGAGAAGAGCTACGAA TTGCCCGACGGGCAAGTAATCACCATTGGCAACGAGAGGTTCCGTGC CCCTGAGGCTATGTTCCAGCCATCCTTCCTTGGTATGGAATCTGCTGG CATCCACGAAACAACTTACAATTCTATCATGAAGTGTGATGTCGACATC AGGAAAGATCTGTATGCAAAACAGTGTCTGCTGGAGGTTCAACTATG TTCCAGGATTGCTGACCGTATGCAGAAAGAGATCTCTGCCCTGGCC CCAGCCACCATGAAATTAAGATCATTGCTCCACCAGAGAGGAAATAC TCTGTATGGATCGGTGGTCTATCTAGCTTCTCTTCCACCTTCCAAC AAATGTGGATTAGCAAGCAAGATATGACGAATCTGGAC**CCTCTATCG TCCATAGGAAGTGC**TCTAA | MDDEVAALVIDNGSGMCKAGFAGD DAPRAVFPSIVGRPRHQGVMVGMG QKDSYVGDEAQSKRGILTLKYPIEH GIVTNWDDMEKIWHHTFYNELRVAP EEHPVLLTEAPLNPKANREKMTQIM FETFNTPAMYVAIQAVLSLYASGRTT GIVMDSGDGVTHTVPIYEGYALPHAI LRLDLAGRDLTDYLMKILTERGYSFT TTAEREIVRDIKEKLCYVALDFDQEM QTAASSSSLEKSYELPDGQVITIGNE RFRAPEAMFQPSFLGMESAGIHETT YNSIMKCDVDIRKDLYANTVLSGGS TMFPGIADRMQKEISALAPATMKIKII APPERYKYSVWIGGSILASLSTFQQM WISKQEYDESGPSIVHRKCF* | GCACTTCCTATGGACGATAG**AGG** | **actin-II_F:**<br>GAAATACTCTGTATG ATCGGTGGTTTC | No detectable cutting (determined by Sanger sequencing) | N/A |
|  |  |  |  |  | **actin-II_R:**<br>GTCTTGTATGTGTAAGCCAGGAAGCTG |  |  |
| *epsilon* / MSTRG.3512 | scaffold1546:2664376-2678998 | ATGGCCGACCGAGATGAAATGGTATACAAGGCTAAACTGGCTGAACAA GCCGAGAGATATGACGAGATGGTGGAAAATATGAAAGAAGTTGCTGAC CAAGGCTCAGAGTTGACTGTTGAAGAAAGAAACCTTTTATCAGTTGCTT ACAAAAATGTGATAGGAGCACGACGAGCTTCGTGGAGGATCCTTAGCA GCGTAGAGCAGAAAGAAGAGCCCAAAGGTTGTGAGCAGCGACTAAAT TTGATTAGATGTTACAGATCAAAGGTTGAGGCTGAGCTACAAGATATTT GCAACGATGTGCTGGCTGTATTGGAGAAAAAACTTTTGGTGAATGCCA CTTCTGGAGAATCTAAAGTGTTTTATTACAAAATGAAGGGCGATTATCA CCGCTACCTGGCTGAATTTGCCACTGGCACTGAAAGGAAGCAAGCTG CTGACAACAGTTTAGCAGCTTACAAGGCAGCCAGTGACATTGCCATGA ATGAGCTGCCACCACTCATCCATCCGATTAGGGCTGGCTCTCAATT TTTCTGTTTTCTACTATGAGATCCTTAGCAACCTGAACGAGCTTGTG GCTGGCCAAAGGCTGCATTTGACGAAGCCATTGCTGAGTTGGACACCTT GAGTGAAGAGAGCTACAAAGATTCCACCTTATAATGAGCTTCTGAG AGACAACCTTACTCTATGGACATCAGATATGCAAGCTGAAGAAATGTGA ACAAAAAAGTGAAGCAGCT**GCAGGATGTGGGAACCGAGGAGG**TGTCAT AA | MADRDEMVYKAKLAEQAERYDEMV ENMKEVADQGSELTVEERNLLSVAY KNVIGARRASWRILSSVEQKEEPKG CEQRLNLIRCYRSKVEAELQDICND VLAVLEKKLLVNATSGESKVFYYKM KGDYHRYLAEFATGTERKQAADNSL AAYKAASDIAMNELPPTHPIRLGLAL NFSVFYYEILSNPERACRLAKAFADE AIAELDTLSEESYKDSLIMQLLRDN LTLWTSMDQAECEQKTEQLQDVG TEEVS* | GCAGGATGTGGGAACCGAG**AGG** | **epsilon_F:**<br>TGAATGTTAGTCTCA CTGTATTAAAGTTAA AACC | 78.53% mutant reads | **15 nt deletion:** AGGATGTGGGAACCG<br>**Result:** 5 amino acid truncation |
|  |  |  |  |  | **epsilon_R:**<br>GTTCTCATATGAGTG TTGTTGTAATGTGC |  | **24 nt deletion:** AGCTGCAGGATGTGGGAACCG AGG<br>**Result:** 8 amino acid truncation |
| *sytt11* /<br>MSTRG.1650 | scaffold124:47276-189626 | ATGGTGCAACAGACTTTGGTTGAAAATCATCCTTTGACTCCACATTCCC<br>TTTCTGCTGCTGCAGTTGTTGGGATATGTTTTGGTGTGCTTGTAATGCT<br>GGGCACAGCATCAGCTATTGCATACAAGTGTTACAAACGACAACTAGC<br>CAAAAATGCTCAGAATCGTCTTTCCTCAGCCATTAAAAAGGGCCTGAAA<br>GAGAGCAACAATGGCAACCTGAAGAAGGTGTCATCAGCTCGAAAAGTC<br>ATCATGAAGTCGAAGGCTACTAGCCCTACATATCCTTCGTCAGGTGAG<br>CACATTAGTGGATCGTTATCTGAACGAGGTCTAGTCAGTGGAAGTGGG<br>GCAGGTCCACTTCCTTCTCAGTTCGATCCCTGCCAGGTAAAAGCAGAA<br>TACTATATTGAGAATGAAAAGGATGGCGCCACTCCTGAAAAAGACGAA<br>AGTATGCGTGAGAAACATGAAGACGAGCTGAAGACACCTCACCTTGG<br>GACTTTGCACTTCTCTACTGAATACGACTCTCAGAAGAGTGCACTTCTG<br>GTCACTATTTTGCAAGCAGCTGAATTGCCTCCTCGTGAACCAAGTGTT<br>GGAGGCTGTGACCCATATATCAAGCTGCAACTTCTACCAGAGAAAAAG<br>CACAAATGCAAAACCAGGGTTCTCAGGAAGACCTTGAATCCTGTGTAT<br>GACGAAACCTTCACGTTTTATGGCATTAGCTTCAACCAGCTGCAGAGT<br>ATCACCCTGCATTTTGTAGTTCTTAGTTTTGACCGATTCTCAAGAGATG<br>AGATCATTGGAGAAGTTATTTACCCCCTGAATGGAGTTGACTTGTCTCA<br>GAAAGAGGTTTCTGTGAGCAAGGACATCACACCAAGGCATCTCAAGAC<br>TGACCTCACTACAACATATATAGAAACTGTGTGTGTAGTAAGCCCAACA<br>TATCCTTCTTCCTGTGAAAATAGAAGATCCCTAACTGAAAGCAATAAAG<br>GGGCTGCTGAACAACACAAATCTCTGCTTGATCCTTGTGATCTTAAGG<br>CTGACTGCTCTGTTGAAAATGAGAAAGAGGATGTGATACAAGAAAAAG<br>ATGAGAGCCAGAGGAACAAACCACGACTGCCCCACCTGGGTACGTTG<br>CACTTCTGGGCCAACTATGATACCCAGAAGAATTCACTCTTTGTGACAA<br>TCCTGGAGGCTGATAACCTTTTGCCAACTGACACAGCTGCAGGCAGCT<br>GTGACCCTTACATTAAACTGCAGCTGTTGCCAGAGAAAAAGTACAAGT<br>GTAAAACTCGAATCCTGAGGAAGACCCTGAATCCTGTGTATGACGAAA<br>CCTTCACATTTTATGGCATCAGCTTCAACCAGCTGCAAAACATCACCCT<br>ACACTTTGTTGTCTTTAGTTTTGACCGATTCTCAAGAGACGAGATCATT<br>GGAGAAGTTATTTACCCACTGAATGGCATTGATTTGTCAGACAAAGAAA<br>TCTCTGTGAGCAAGGACATCACACCAAGATATCTCAAGTTTCGTACCC<br>AGGGACGGGGAGAGTTGCTGGTTTCTCTTTGTTATCAGCCTGCTGCTA<br>ACAGACTTACAGTTGTTGTACTCAAAGCAAGGAATCTTCCTAAAATGGA<br>TGTCACTGGACTTTCTGATCCCTATGTGAAGATTTACCTTCTGTACAAT<br>GGGCAGAGAATTGCAAAAAAGAAAACACATGTGAAGAAACGCACCTTG<br>AATCCTGTGTTCAATGAATCATTCCTGTTTGATGTTCCTTATAATGAGG<br>GTCTACAGAATATCAGCCTTGAGTTCCTGCTTCTTGATTATGACCGAAT<br>GACAAAGAATGAAGTCATTGGCAGATTGGAAGTGGGTGCTCGCACTTT<br>GGGTCAAGAGTTGCATCATTGGAATGAAGTAATGAACTGTCCACGTAA<br>GCAGATTGCTGAATGGCACAAACTGAAAGAATAA | VQQTLVENHPLTPHSLSAAAVVGIC<br>FGVLVMLGTASAIAYKCYKRQLAKN<br>AQNRLSSAIKKGLKESNNGNLKKVS<br>SARKVIMKSKATSPTYPSSGEHISGS<br>LSERGLVSGSGAGPLPSQFDPCQV<br>KAEYYIENEKDGATPEKDESMREKH<br>EDELKTPHLGTLHFSTEYDSQKSAL<br>LVTILQAAELPPREPSVGGCDPYIKL<br>QLLPEKKHKCKTRVLRKTLNPVYDE<br>TFTFYGISFNQLQSITLHFVVLSFDRF<br>SRDEIIGEVIYPLNGVDLSQKEVSVS<br>KDITPRHLKTDLTTTYIETVCVVSPTY<br>PSSCENRRSLTESNKGAAEQHKSLL<br>DPCDLKADCSVENEKEDVIQEKDES<br>QRNKPRLPHLGTLHFWANYDTQKN<br>SLFVTILEADNLLPTDTAAGSCDPYIK<br>LQLLPEKKYKCKTRILRKTLNPVYDE<br>TFTFYGISFNQLQNITLHFVVFSFDR<br>FSRDEIIGEVIYPLNGIDLSDKEISVSK<br>DITPRYLKFRTQGRGELLVSLCYQP<br>AANRLTVVVLKARNLPKMDVTGLSD<br>PYVKIYLLYNGQRIAKKKTHVKKRTL<br>NPVFNESFLFDVPYNEGLQNISLEFL<br>LLDYDRMTKNEVIGRLEVGARTLGQ<br>ELHHWNEVMNCPRKQIAEWHKLKE<br>* | N/A | N/A | N/A | N/A |

**Table S1. Gene sequences, CRISPR gRNAs, and genotyping primers.**

|  | Optic lobe (OL) | Posterior sub-<br>esophageal<br>mass (PSM) | Vertical lobe | Stellate<br>ganglion (SG) | Retina | Sucker | Tentacle | Skin |
| --- | --- | --- | --- | --- | --- | --- | --- | --- |
| *actin I* / MSTRG.17104 | 50.9 | 50.7 | 26.6 | 100.2 | 96.3 | 3915.5 | 7439.1 | 201.8 |
| *actin II* / MSTRG.58 | 1883.0 | 2633.6 | 1788.5 | 3173.7 | 2877.0 | 2019.3 | 1675.4 | 2650.7 |
| *epsilon* / MSTRG.3512 | 170.8 | 450.3 | 327.4 | 429.0 | 109.7 | 91.4 | 79.7 | 145.6 |
| *syt11* / MSTRG.1650 | 171.6 | 179.2 | 133.1 | 165.9 | 131.1 | 2.0 | 4.0 | 0.7 |

**Table S2. RNA-seq FPKM values across tissues.**

| Gene | ATAC-seq defined regulatory element |
| --- | --- |
| **actin II /**<br>MSTRG.58 | AGAAACCAATCCTTTCTAAAGCAAAGTGGTTGTTTTAGAGACCAACTATTTACTGGCCAACTTATTGCAATTGAGATTGTTTGCATTGCTTCTTCTCTATTTGGTGAAAAATAAACCCCTCTCTCTT<br>TACAACTTACTCTTTCGTTCATTGATCATTGTCTTACACTCACAATCTCTTTTTTTTTAGTTCTTTCTAGTTTCTTTTTATTTATTTTTTATCTCACTCCGCGTCTTTTCTTACTCAACCTTTCTCCTTC<br>CCGAGTTTATTGTTATGCGACCACTTTTTTGCTTTCTATCTCTCCTGTCGCTCTCAATGCCTTTTCTCTCAGTTTACTCGTTTCTTCTTCCCTGATCAGTCTTTTTTTCCCCACCCATCCTTTCTTC<br>CTTGTTATGCATTGTTTTTCTTGGCTTTCTCCTGACGCCACTCCTCTCCTTCTCGTAAATCCCGCCTCCTCCTCTCTCATTACCTTTATGAAATCATCATCATCGCAGTAAGGCTAACAGTAGCAC<br>GTCTTAAATGGCATGCTTGCTTTGACAGGAAAGACGCGCCAATCGTGACCATATATGGTAATAGTACCACTTCCATATAAAGTTAAACTCTTCATATGAGAAATATTTTTCCTGGAACTCTTTCGT<br>CTTTCTTACAAAATAGGCTGTTAAGCTATTGTGTGTGCGGCAGTATTTAAAGCGGGCAGACACCAATTTCCACAGTCTAGTATTTCACTCCACCGGACACACATCTCGGTTGGTCTGCATTCACT<br>TCGTCTGAAGACATTTGAACAAGGCAGTTGTCACTTGCCAATTTTAAATTTTTTCTACTAAAATCTTCTGTTGTTGTAGTTTTGACTACCTTTAAACCCTACAAATCGTCGTTATTCAAGTTGCTGC<br>TCTCGTTATTGACAACGGATCTGGTGTCTGCAAAGCTGGTTTTGCCGGTGATGATGCACCTAGGGCTGTATTCCCATCTATTGTCGGCCGTCCCCGACATCAGGTAAGAATACTTTATTTTATTT<br>TATTATATTTATTTTACCGTTCAGAAAGCCTTTATTTCTTAGTTTTGACACTTATCTCTTAAGGCTTCCTTTCCTGTAATTTTAAAATTTTTAACTTTTTATCTTGTTTCTTGAAATAGCACAATTATTC<br>CAATCCGTAAGAGTTCAAGCTTTTTTTTTTTTTTTATTTGATTCTTGATACTGTAATTTTTTACTAAAAAAAAAAAAAAAAAAAGTTAATGTTTGCTTCAATATGCTTATTAAACAATGCTATCTTTATT<br>CACTTCGGTTTACTTTCTCTTACCGTAACTTGAATAACCATGAACTGAAAATTTAGCAAGCATGTTTTCGAGCTGGTTACGACAGCCTTGAAAATATAGCATGATTTTGCAGTTTGGCTGGCTATA<br>ACCTTATATGGTATTGCACTACCATATATGGTCATTACCATGAATGGTAGAAAATGCCAGTCTGCATTCTGATTGGCTCTTAAAAGCTATCCTAGGGAACGGCAGCGCTCTGCATATTTTTGAAG<br>CGTTGGCTGACTCATTGAAAGGGGGTGGGAACTTAAAAGGCCGGTTTCTTAACTTAAACGGTAATCAATTTGACATTATTTGTGCTGTTTTAATATTTACTGTAATTTTTTTTTTCTTTAATTTTGT<br>TTAAAACTTCACTTTTAAATTTCGTTTTCTGCAATCATTTTAAGCTTGTTATTAGGGTTCTACGTCACTCGGCTCTAAGTCTACCATCCACTTCCAAATATGGTCACAGAAAGTTCTGTAGGCGTG<br>GCATTTTTTATGATGGGTAGGCACGTGATAATGGACTTGGCGTAAAATATTTCAATATTTTAACGTGATCAACCCTCTAAAAATTTAATTACTTTTTACAAGAATTAACTGATGCTTGTTTATTTGA<br>ACTTGCCAAGTACCTGTCGCAAGTGGCTGTATTTTACTCTTTATCGGCATATTTAGATTTTTATTCCCCCCCCCTCCCCCCAACAGTAACGAAACTTTCACTGCTTTTTGTGAAAGCAACTTCAGT<br>TTGGTGCCCCAAATAAATTATCACGATTGATACGGCACTGACAGTCGCGAAAACTACTTGTTTTCATTTATTTCTGTTAACAGTATTGGTAATACTTCAGAAATGCATTTTTTTTCAAAACGAAGCT<br>TGCATGCTCTGAATTTTGACCGTTAAAAAATGTTATGAAGCTCATTGATTTGTTTCATTTTAAAGTACTTTTGTGTTTTAAAAAAGCTTCTTCTCCTCCCCCCCCCGCCCATGCCTCTTTATACAAAATACTAAGTCTTTTTTA<br>CTCTAAATTTTTTTAACTTCGTAACATATAGCCTTTATTAAGTTGGGACGCTAGACAATAAATGAATTTTTGTTTCTCTCCCCCCCCCGCCCATGCCTCTTTATACAAAATACTAAGTCTTTTTTA<br>TTTCTATCGTTCAGGGG |
| **synt11 /**<br>MSTRG.1650 | GTAAATGATGTAGCAAGAGAAAACGTGTTTTAAGTGA AAAACAAATAAAGTAGAAGGAGGAATTTAGAAAAGTGGACCTACAAACGATGAATATGAAGTGAGAATAAGCAAAATGGAGTTTGAA<br>GAGTAATATAAGAAGAAAAGAAATATTGATTTTGTTGTTCAACTGAGGTTGTTGCTGCCACTGCTGGGCTTTTTCTACTGCAGCCTCTTCGCCATAATATGATGCAGTTTCATTCAGAGACAGTT<br>CAATATCATTATCAGTGTTATTAGTGGTGCTGTTGCTGAATCGTATTAGTAACGAATAGGGTAAGGGAGATTCAAATCAGAGAGCAAGGCGAACCTTACTCTCCTCTTACAGATCGATACGCAA<br>CAGACTCGGCCATCAACAGCAGTAGCAGTTGTAGATTCGTTGTTGATTTGCCTTCTCTCTTAGCTCACTCATCAATCTCTACCTCACTCTAATAATTCTGCCTGTCAACATTATCCCCTCTTTCAC<br>CACCTCCTTATTTATCAATATTAACAATTTTACTTCATCCATTTAAATCATACCTTAAACTCTTTCTTCACTCTTTGTTTGTCTCATTTAGCTAACCTTTTCTCATCTTTTTCCTTTTCCTTATCTTCCT<br>CTTACTCCTCACATCATAATACTGTTTCTTACTTCAGTTCTCTATAGGGCGTTGCACCTCTTTCCCCCTTATCTGCATACTTATCATCCTTTTCTCTCAATATCTCTCTCCCTCACTTTGTACATCC<br>ATCTCGTATCCATCCTATTTTATTGTCTCCTCCACCTGTACGTTTACCTCTGACTACATCTCATTTTCCCTCTCTCTTTTAAAGTATACTTAATCTTGCTTTTCCCCTTTGTCTAGTTTACTCGCACT<br>TGACTCTCCGGCGCCCTCTTTGCTATTTGTTTTCCCACTCTCTCATTCATCCTTATCTCTTTTTCCCCTTCCTTGTCGCATCTTCATCATTGGAGAAGGTGATTGAATAATATCATCATCGCCAAC<br>GACAAGGGACAAAGAATTTCCCGGCTCACAAGCTTCTTTCTGCCATTCTTCTTCGGTTTTAGACAGTAGTAGTGATTACTCTCTACTGCTACTTTTTTTTTTTTGAAATATCTGATTCTAATTTATA<br>GGCCAAGACAAACAAATAATCCACATACACCTCTACATACATACATATTTACACAAATACACACATACACGCAC |

**Table S3. Sequence of ATAC-seq defined elements from *actin II* and *synaptotagmin-11* used in transgene constructs.**

|  | Ctx1 | % | Ctx2 | % |
| --- | --- | --- | --- | --- |
| **Embryos injected** | 106 |  | 43 |  |
| **Survival to hatching** | 1 | 0.94% | 0 | 0.00% |

**Table S4. Low survival following CRISPR targeting of cephalotoxin genes.** *cephalotoxin 1* (*ctx1*) and *cephalotoxin 2* (*ctx2*) are venom genes expressed in the posterior salivary glands of hatchling and adult cuttlefish<sup>2</sup>. Given their predicted roles in prey envenomation, these genes are not expected to be required for embryonic development. Nevertheless, survival to hatching following Cas9/gRNA injection targeting either gene was low.

### References

1. Montague, T.G., Rieth, I.J., and Axel, R. (2021). Embryonic development of the camouflaging dwarf cuttlefish, *Sepia bandensis*. *Dev Dyn*, 250, 1688–1703. 10.1002/dvdy.375.
2. Naidu, M.P., Pardos-Blas, J.R., Attarde, S., Achimba, F., Hempel, B.-F., Clotea, I., Stambouli, B., Kirchhoff, K.N., Williams, M., McCarthy-Taylor, J., et al. (2026). Lineage-Specific Venom Gene Expression Shapes Chemical Diversity in Cephalopods. *bioRxiv*, 2026.04.09.716377. 10.64898/2026.04.09.716377.
